# Identification of novel HPFH-like mutations by CRISPR base editing that elevates the expression of fetal hemoglobin

**DOI:** 10.1101/2020.06.30.178715

**Authors:** Nithin Sam Ravi, Beeke Wienert, Stacia K. Wyman, Jonathan Vu, Aswin Anand Pai, Poonkuzhali Balasubramanian, Yukio Nakamura, Ryo Kurita, Srujan Marepally, Saravanabhavan Thangavel, Shaji R. Velayudhan, Alok Srivastava, Mark A. DeWitt, Jacob E. Corn, Kumarasamypet M. Mohankumar

## Abstract

Switching hemoglobin synthesis from defective adult beta-globin to fetal gamma-globin is an effective strategy for the treatment of beta-hemoglobinopathies. Fetal hemoglobin expression is down-regulated in the postnatal period due to the interplay of transcription regulators with the *HBG* promoters. However, in the hereditary persistence of fetal hemoglobin (HPFH) condition, naturally occurring point mutations in the *HBG* promoter causes continued expression of fetal globin even during adulthood. Inspired by this natural phenomenon, we screened the proximal promoter of human *HBG* genes using adenine and cytosine base editors to identify other nucleotide substitutions that could potentially lead to elevated levels of fetal globin. Both the base editors efficiently and precisely edited at the target sites with a minimal generation of indels and no deletion of one of the duplicated *HBG* genes. Through systematic tiling across *the HBG* proximal promoter, we identified multiple novel target sites that resulted in a significant increase in fetal globin levels. Further, we individually validated the top eight potential target sites from both the base editors and observed robust elevation in the fetal globin levels up to 47 %, without any detrimental effects on erythroid differentiation. Our screening strategy resulted in the identification of multiple novel point mutations and also validated the known non-deletional HPFH mutations that could elevate the fetal globin expression at therapeutically relevant levels. Overall, our findings shed light on so far unknown regulatory elements within the *HBG* promoter that normally mediates fetal globin silencing and identify additional targets for therapeutic upregulation of fetal hemoglobin.

## Background

Fetal hemoglobin (HbF) is a tetramer consisting of two alpha-globin chains and two gamma-globin chains, which expresses profoundly during the fetal stage of human life. The expression of fetal hemoglobin is then silenced progressively after birth until it constitutes only about 1% of total hemoglobin (1). Naturally occurring mutations in the regulatory regions of the gamma-globin (*HBG*) gene have shown to reactivate expression and increase HbF levels during adult life (2). This inherited genetic condition is benign and is known as the hereditary persistence of fetal hemoglobin (HPFH). Individuals, who inherit HPFH mutations along with genetic disorders affecting the adult beta-globin gene, such as sickle cell disease or β-thalassemia, were shown to have milder forms of the disease (2,3). Hence, high levels of HbF expression are considered beneficial for improving the clinical outcomes of patients with sickle cell anemia and β-thalassemia.

Genome editing approaches have largely focused on the beneficial effects of HPFH mutations to increase HbF levels in sickle cell disease (4). These mutations either create *de novo* binding sites for erythroid activators or disrupt the binding sites of repressors, thereby increasing the expression of HbF. For example, the -175 T>C, -198 T>C and -113 A>G HPFH point mutations create *de novo* binding sites for the erythroid master regulators TAL1, KLF1, and GATA1 respectively (5–7). Similarly, the introduction of HPFH associated mutations around -115bp from the transcription start site (TSS) of *HBG*,–114C>A, – 117G>A, and a 13 bp deletion (Δ13bp), and around -200bp from the TSS, –195C>G, - 196C>T, -197C>T, -201C>T and -202 C>T/G, were shown to disrupt the binding sites of the two major fetal globin repressors, BCL11A and ZBTB7A, respectively (8). However, the roles and locations of other key regulatory elements in the *HBG* promoter are relatively unknown. Thus, tiling the *HBG* promoter using base editors could unravel molecular mechanisms of human hemoglobin switching and reveal additional point mutations that could be useful for therapeutic gamma-globin de-repression.

Targeted introduction of HPFH mutations into the *HBG* promoter by nuclease-mediated homology-directed repair (HDR) is relatively inefficient. It often results in high rates of random insertions and deletions (indels) through non-homologous end-joining (NHEJ) DNA repair pathways (9), and in some studies, HDR-generated alleles are not maintained *in vivo* (10,11). Besides, due to the high homology between the duplicated *HBG1* and *HBG2* genes, simultaneous editing of both *HBG* promoters by programmable nucleases that cause double-stranded breaks (DSBs) frequently results in deletion of the *HBG2* gene including the ∼4.9 kb *HBG* intergenic region (12). This deletion would make the *HBG1* gene solely responsible for fetal globin expression in the edited cells and could reduce the overall expression of therapeutic HbF (12).

To overcome these limitations, we implemented a strategy to screen and identify potential regulatory mutations within the proximal promoters of the two human fetal globin genes *HBG1* and *HBG2.* We employed CRISPR base editing to introduce an array of point mutations into the *HBG* promoters and then screen for those mutations that induce HbF to therapeutic levels, without any confounding effects of creating DSBs. We found that base editing using adenine and cytosine base editors (ABEs and CBEs, respectively) is highly efficient in creating point mutations without inducing DSB breaks (13,14). We identified several novel point mutations that are associated with a significant increase in gamma-globin expression and could be of potential therapeutic interest. Our results provide proof of concept for the use of CRISPR base editing to reactivate the expression of gamma-globin for the treatment of beta-hemoglobinopathies.

## RESULTS

Previous studies have shown that the highly homologous *HBG1 and HBG2* proximal promoters play a crucial role in the gamma-globin expression and contain many of the key HPFH associated point mutations (15). To identify novel regulatory elements in the human *HBG* promoter that influence gamma-globin expression, we performed a base editing screen to introduce point mutations in all compatible locations within 320 bp upstream of the TSS of the *HBG* gene (**Fig. 1a**).

**Figure 1:**
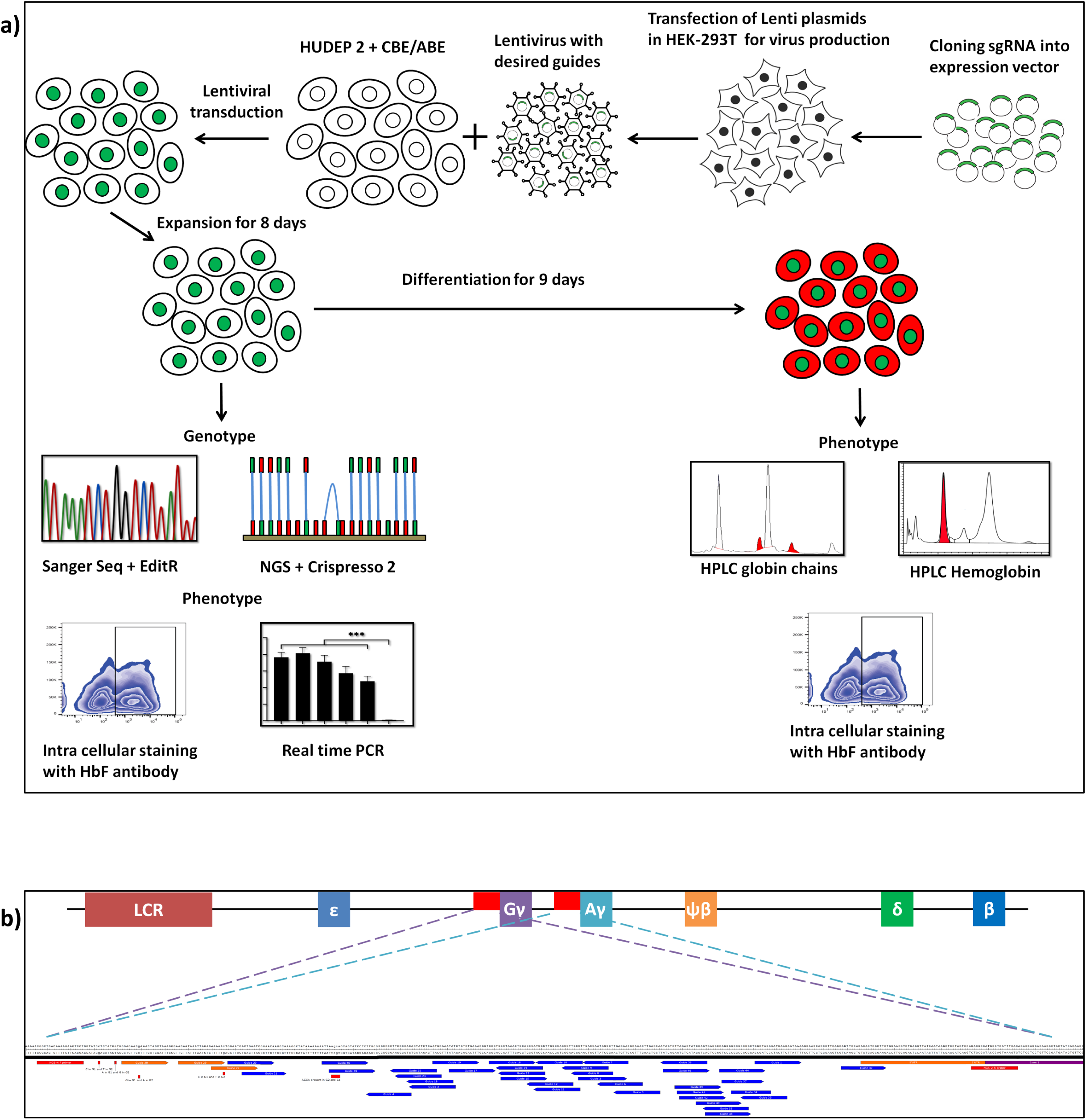
Overview of screening of *HBG* promoter using base editors to identify novel point mutations to elevate HbF: (a) Schematic representation of the overall screening approach, ABE or CBE expressing HUDEP-2 cells were transduced with gRNAs that target the proximal promoter of the *HBG* gene. The edited cells were expanded for eight days. Editing efficiency was evaluated by Sanger sequencing and NGS, while functional analysis was carried out using FACS and qRT-PCR. Top targets from both the ABE &CBE screens with the highest induction of HbF were validated and differentiated to erythroid cells. The differentiated cells were further subjected to FACS, RP-HPLC, and HPLC analysis to determine the number of HbF positive cells, individual gamma-globin chains, and fetal hemoglobin levels, respectively. (b) Representation of gRNA targeting *HBG* promoter region in HUDEP-2 cell line, gRNAs targeting -320bp upstream of TSS in *HBG* genes (*HBG1* and *HBG2*) promoter regions are represented in the figure. gRNAs common for *HBG1* and *HBG2* promoters are represented in blue, while the gRNAs specific to *HBG1* promoter is represented in orange color, the primers used for deep sequencing are represented as a red bar.

HUDEP-2 cells, an immortalized human erythroid progenitor cell line (16), were transduced with ABE or CBE lentiviral constructs to establish stable cell lines which expresses either ABE or CBE (HUDEP-2-ABE or -CBE). After stable cell line preparation, gRNAs targeting the *HBG1 and HBG2* proximal promoter were designed, and gRNAs with a suitable base editing window (target nucleotide in position 3-9 from PAM distal end) for ABE and CBE were chosen (**Fig. 1b**). Among the 41 gRNAs designed, 36 gRNAs had a base editing window for ABE editing, and 32 gRNAs had a base editing window for CBE editing (**Sup. Table 1**). We transduced HUDEP-2-ABE or -CBE cells with the respective gRNAs and expanded the cells for eight days. Successful base editing was then confirmed by NGS analysis of the target sequences (17).

First, we determined the overall efficiency of ABE-induced A to G conversions and CBE-induced C to T conversions at different target sites of *HBG* promoter and found that the base editing efficiency varied from 0-82% and 0-78%, respectively (**Fig. S1**). In case of ABE, of the 36 gRNAs that were screened, seven gRNAs showed very low base editing (<10%), and most of the gRNAs (29 gRNAs) showed base editing efficiency between 10% to 82%. In case of CBE, out of 32 gRNAs that we screened, around 12 gRNAs showed a low base editing rate (<10%), and most of the gRNAs (20 gRNAs) showed base editing efficiency between 10% to 78% (**Fig. S1**). The individual base conversion of A to G was observed at the on-target site of all 36 gRNAs with the base editing frequency ranging from 0-74% for ABE; similarly, the individual base conversion of C to T was observed at the on-target site of all 32 gRNAs with the base editing frequency ranging from 0-61% for CBE (**Fig. 2**).

**Figure 2:**
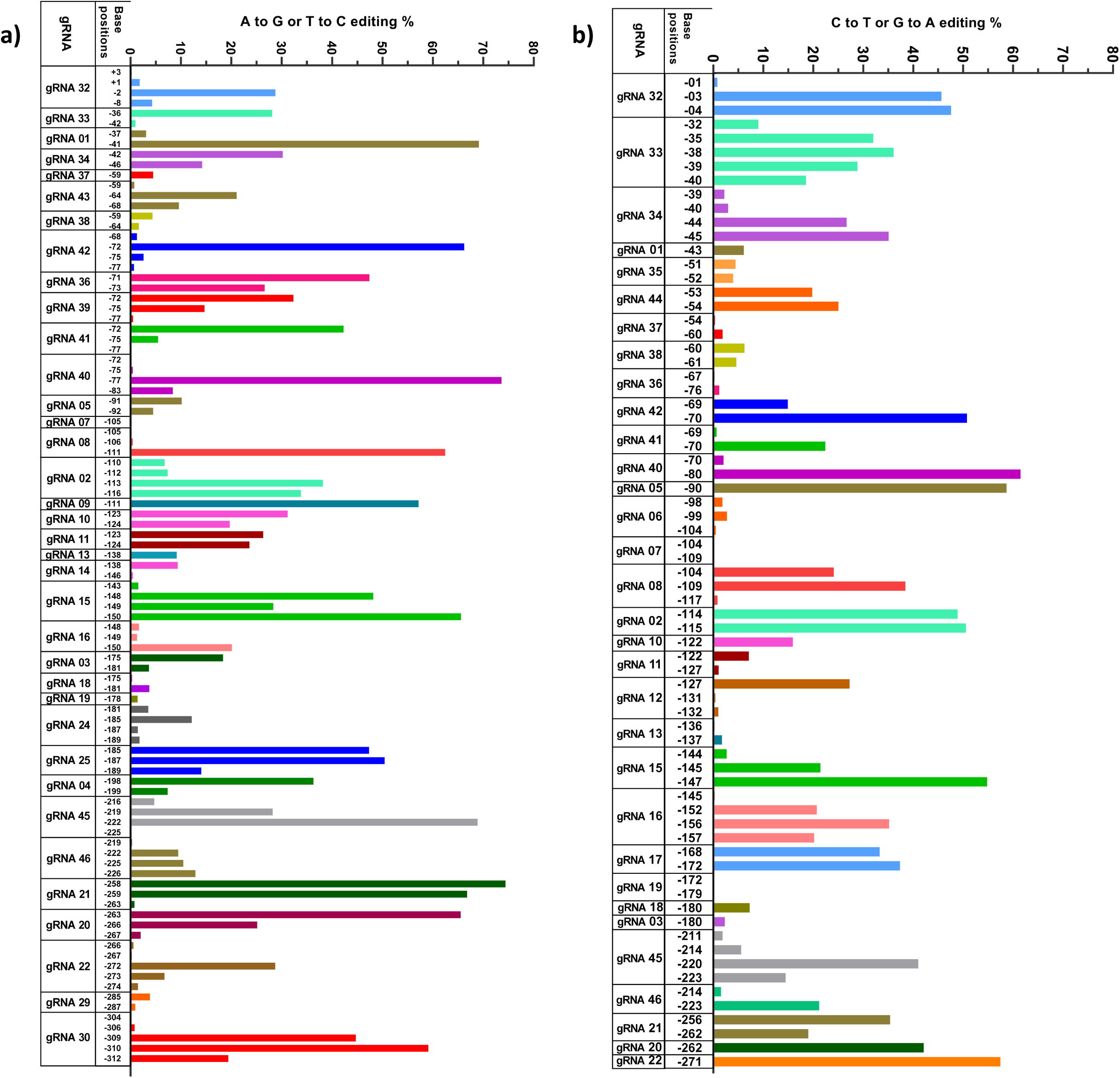
Analysis of base substitution efficiency in *HBG* promoter by ABE and CBE through NGS: HUDEP-2 cells expressing ABE (a) or CBE (b) transduced with different gRNAs targeting the proximal promoter of *HBG* were deep sequenced by NGS and analyzed using Crispresso-2. The base conversion positions are sequentially represented in the x-axis up to -320 bp from the TSS. The base conversion efficiency of individual bases for each gRNA is represented in the y-axis as the percentage of the desired base converted within the respective read (A to G or C to T).

The base substitution efficiency for each target varied significantly depending on the base editing window for ABE and CBE. The average editing efficiency observed was high (>30%) in case of canonical position for ABE (A5-A7) and CBE (C5-C7), while it varied between 1-27% in the non-canonical position for ABE (A1-A4, A8-A12) and CBE (C1-C4, C8-C16) for the different gRNAs used in this study. (**Fig. S2**).

Subsequently, we determined the effect of ABE- and CBE-mediated base editing on the product purity and indel frequency for all gRNAs. Most of the gRNAs in CBE transduced cells showed unanticipated C- to non-T edits (C-R/G-Y), among which C to G conversion was predominant. In the case of ABE, we observed minimal level of targeted bases getting converted to unexpected bases (A-Y/T-R) at a few on-target sites, consistent with previous studies (13,14) (**Fig. S3**). ABE and CBE stables transduced with respective gRNAs showed less than 2% of indels when analyzed using deep-sequencing data (**Fig. S4**). In agreement with our NGS results, Sanger sequencing (18) further confirmed the base substitution efficiency at the target loci (**Fig. S5).** In summary, our results suggest that ABEs exhibit higher product purity and lower indel frequency than CBE in all cases.

To evaluate whether the targeted base substitution at the *HBG* promoter using ABE or CBE has increased HbF expression, we analyzed the edited cells by flow-cytometry after HbF staining. The percentage of HbF positive cells ranged from 3.1%-48.6% in ABE and 1.1%-39.2% in CBE (**Fig. 3**). In our preliminary analysis, among the 36 gRNAs that we screened for editing with ABE, five gRNAs showed a high increase in the number of HbF positive cells (in a range of 40% to 50%), twenty-two gRNAs showed a moderate increase in the number of HbF positive cells (in a range of 10% to 30%) and nine gRNAs resulted in fewer HbF positive cells (in a range of 3.0% to 10%) as compared to control (4.65%). In case of CBE, one gRNA out of 32 gRNAs showed a very high increase in the number of HbF positive cells (about 40%), 12 gRNAs showed a moderate increase in the number of HbF positive cells (in the range of 10 % to 18%) and 19 gRNAs resulted in fewer HbF positive cells (in a range of 1.0% to 9%) compared to control (4.4%). Interestingly, in both ABE and CBE experiments, gRNA-2, gRNA-10, gRNA-11, and gRNA-34 were the top eight targets that showed an increase in the percentage of HbF positive cells. In summary, the primary screening of the *HBG* promoter by ABE and CBE revealed that only a few gRNA targets are important for the *HBG* regulation.

**Figure 3:**
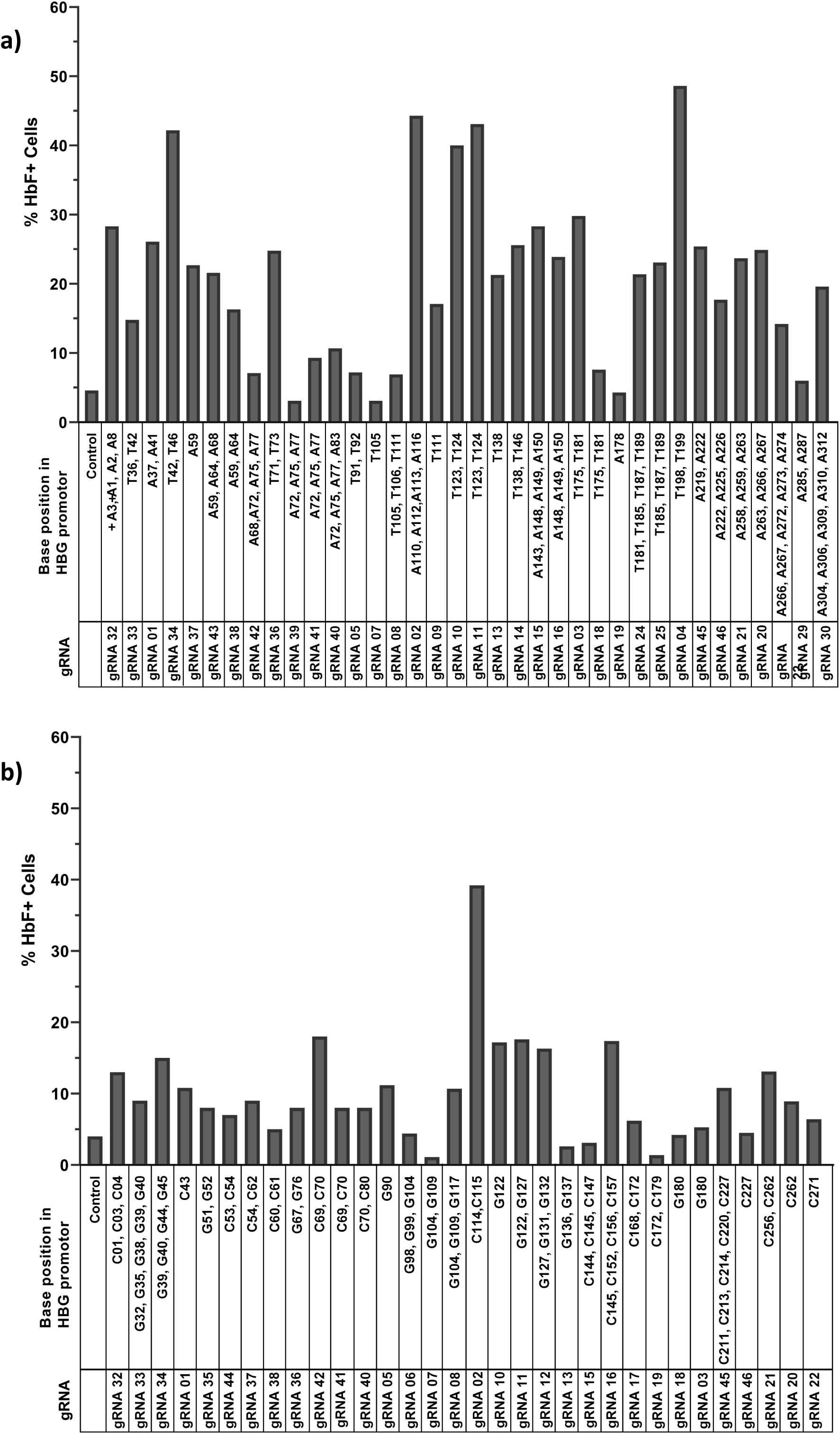
Base editing of the *HBG* promoter reactivates developmentally silenced gamma-globin expression in HUDEP-2 cells. Percentage of HbF positive cells in ABE (a) or CBE (b) expressing HUDEP-2 cells transduced with the respective gRNAs analyzed by flow cytometry. The edited base positions in the *HBG* promoter region for the respective gRNAs are represented in the x-axis.

We further validated the top eight gRNAs -2, 3, 4, 10, 11, 15, 32 and 34 from the ABE screen and gRNAs -2, 10, 11, 16, 21, 32, 34 and 42 from the CBE screen which gave the highest level of HbF positive cells. The top-scoring gRNAs identified from this screen includes gRNAs-2, 3, and 4 that recreated the well-known HPFH mutations -114C>T, -117G>A, −175T>C and -198T>C (5,8,19,20) and additional gRNAs that are potential new targets. The total editing efficiency of ABE (ranged from 22-77%) and CBE (ranged from 12-83%) and the individual base conversion of A to G (ranged from 22-77%) or C to T (ranged from 22-77%) at the respective target regions were comparable with the screening results. Undesired modifications were observed in CBE but not in ABE (**Fig. 4 a-b**), and the overall indel percentage in both ABE and CBE was less than 2% (**Fig. S6**).

**Figure 4:**
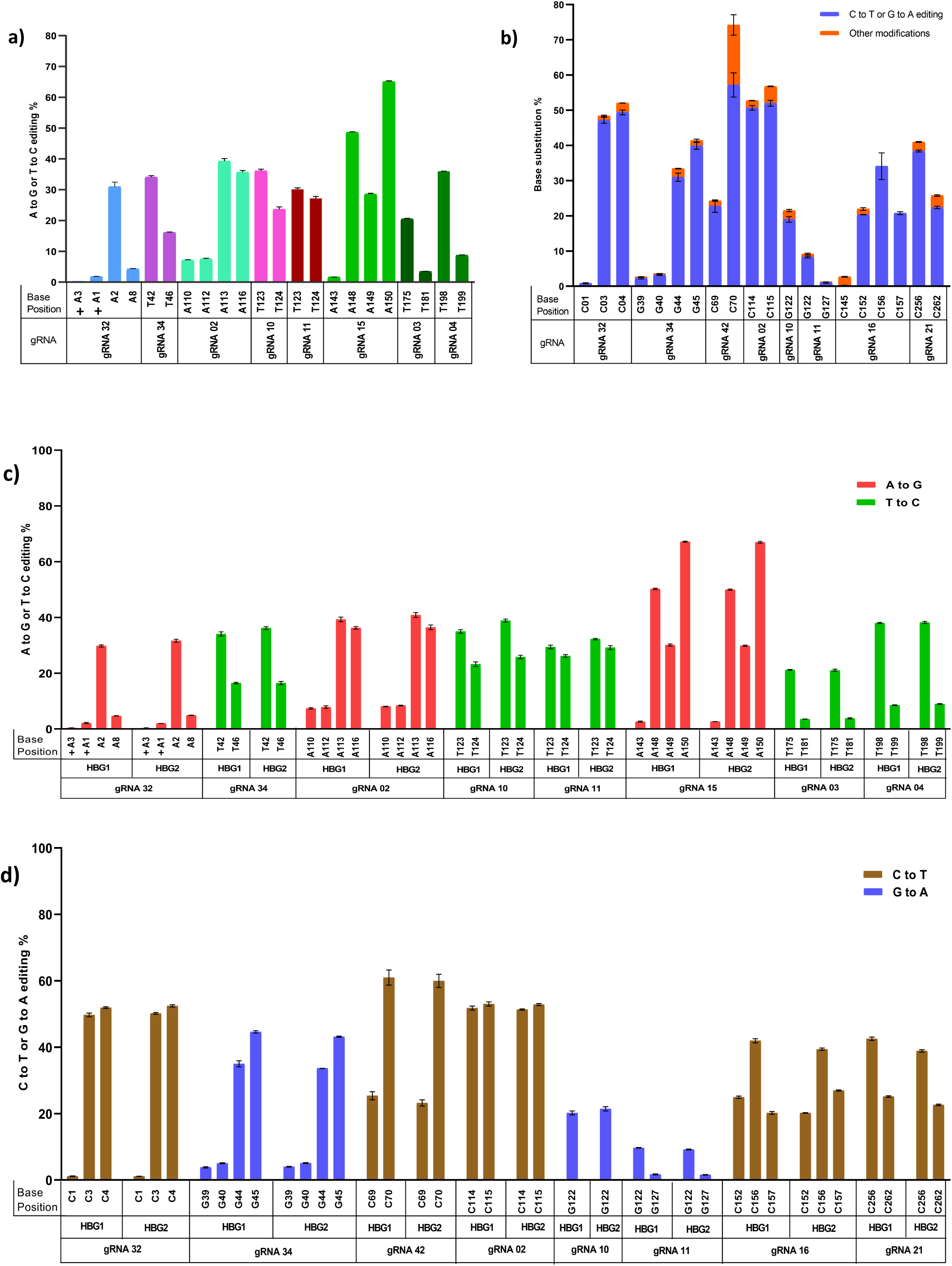
Validation of targeted base editing for top eight gRNAs from the primary screen of ABE and CBE at *HBG1* and *HBG2* promoters. HUDEP-2 cells expressing ABE or CBE transduced with the top eight gRNAs were analyzed by deep sequencing at the targeted regions in the *HBG* promoter. The editing efficiencies of ABE (a) or CBE (b) are represented as the percentage of total sequencing reads with target C:A converted to T:G at specified sites. The edited base positions in the *HBG* promoter region for the respective gRNAs are represented in the x-axis. Comparison of base editing efficiencies of ABE (c) or CBE (d) at the indicated target sites of *HBG1* and *HBG2* promoter in HUDEP-2 cells. The highly homologous *HBG1* and *HBG2* promoter regions were amplified and deep sequenced to segregate the specific editing between the promoters by using single nucleotide variation at positions -271, -307, -317 and -324 between *HBG1* and *G2* by using Bowtie and analyzed through IGV software. The editing efficiencies are represented as the percentage of total sequencing reads with target C to T conversion or A to G conversion at specified sites. Data are expressed as mean ± SEM from three biological replicates (p>0.05).

To find out whether there is any variation in base substitution efficiency between the highly homologous *HBG1* and *HBG2* promoters, we performed deep sequencing using a standard primer, which amplifies both the *HBG* promoters. We determined the specific editing in *HBG*1 and *HBG*2 promoter based on single nucleotide variation at positions -271, -307, -317, and -324 by using Bowtie and IGV software (21,22). Our analysis showed that base editing rates were highly similar in the *HBG1* and *HBG2* promoters (**Fig. 4 c-d**).

After validating the editing efficiency, *HBG* expression was determined using quantitative real-time PCR. We observed a significant increase in the *HBG* mRNA expression for all the top eight gRNAs in ABE edited cells (p<0.001). In the case of CBE, gRNAs 2, 10, 11, and 42 showed a substantial increase in *HBG* mRNA expression (p<0.01), while gRNAs 16, 21, 32, and 34 showed a modest level of expression (p<0.1) as compared with the control (**Fig. S7 a-b**). Next, we analyzed the level of HbF positive cells by flow cytometry in the ABE and CBE edited cells before differentiation. HbF positive cells in ABE ranged from 24-50%, and in the case of CBE from 11-60%, while in control, it was around 0.8% and 3.9 %, respectively (**Fig. 5 a-b and S9 a-b**). The percentage of HbF expressing cells was significantly increased in both ABE and CBE edited cells compared to the control, which was consistent with the qRT-PCR results.

**Figure 5:**
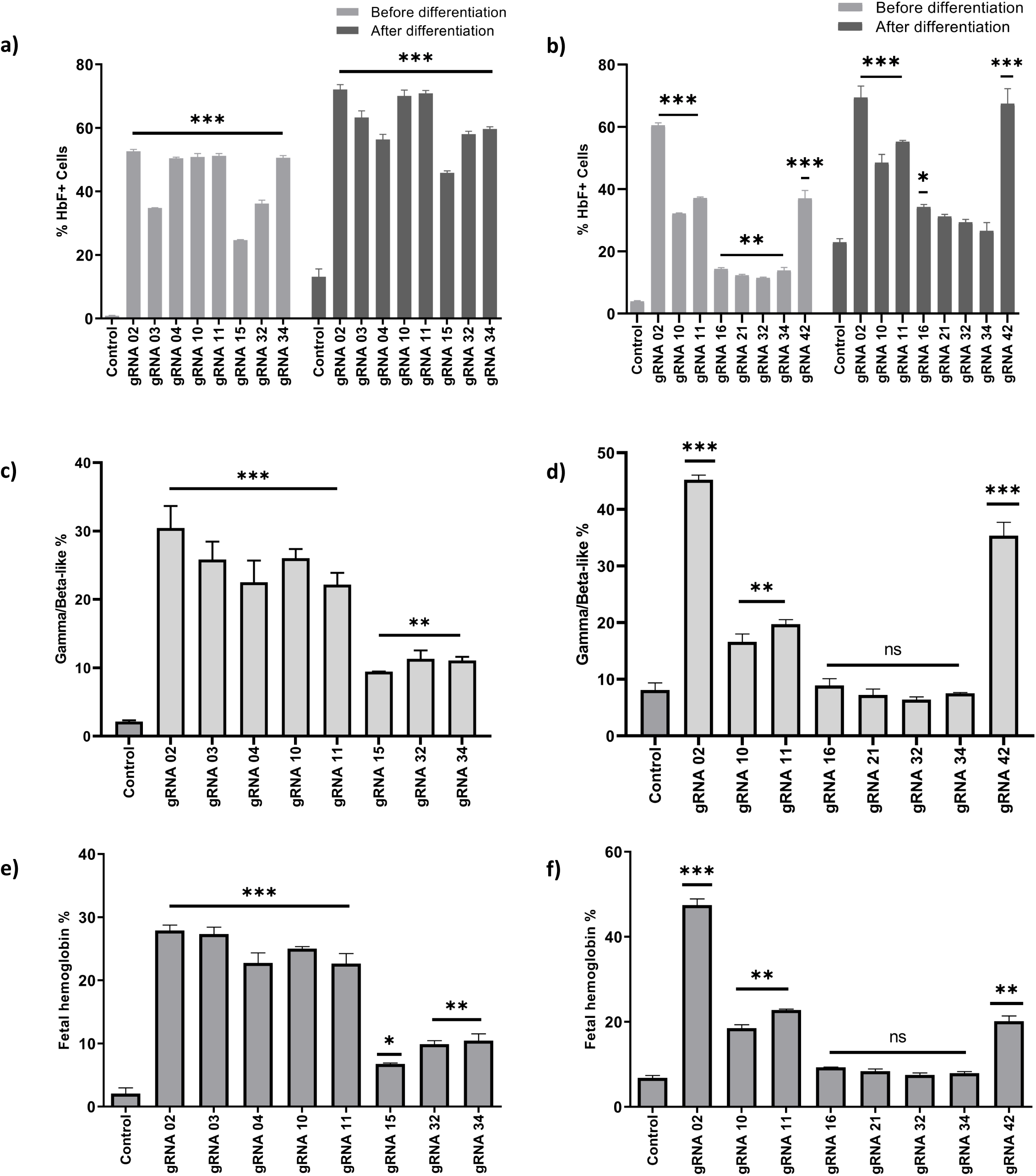
Upregulation of HbF upon base editing on the potential target sites in *the HBG* promoter. Evaluation of HbF positive cells in HUDEP-2 cells expressing ABE (a) or CBE (b) transduced with the respective gRNAs, before and after differentiation by flow cytometry; Globin chains analysis in ABE (c) or CBE (d) edited HUDEP-2 cells after erythroid differentiation by RP-HPLC; Fetal hemoglobin analysis in ABE (e) or CBE (f) edited HUDEP-2 cells after erythroid differentiation by HPLC. All data are expressed as mean ± SEM from three independent biological replicates. Asterisks indicate levels of statistical significance ***p<0.001; **p<0.01

We then determined the effect of base editing on erythroid differentiation by using flow cytometry analysis of CD235 and CD71 markers. The shift in expression of CD71 positive cells alone to CD71/ CD235 double-positive cells determines the erythroid differentiation pattern of HUDEP-2 cells (16). The percentage of double-positive cells was around 83-90% for CBE and above 95% for ABE edited cells when compared to the control, which was 77% and 97%, respectively, suggesting that the differentiation ability of the cells were not affected (**Fig. S8 a-b**). Along with the differentiation markers, the edited cells were stained with the HbF antibody to evaluate the percentage of HbF positive cells. As expected, the percentage of HbF positive cells in differentiated erythroid cells were slightly higher than that of the undifferentiated cells (**Fig. 5 a-b**).

Finally, the differentiated erythroid cells were further analyzed using HPLC to determine the level of HbF production (**Fig. S8 c-d**). We observed a significant induction of HbF in all the ABE edited samples (gRNAs 2, 3, 4, 10, 11, 15, 32, and 34) (**Figs. 5 c, S8-c, and S9 e**). The percentage of HbF was around 6-47% in CBE edited samples, among which gRNAs 2, 10, 11, 42 resulted in HbF above 15%, which is significantly higher than the control (**Figs. 5 d, S8 d and S9 f**). Similar results in terms of elevation of gamma-globin chain levels were obtained in both the ABE and CBE edited samples when compared to the control (**Figs. 5 e-f, S7 c-d and S9 c-d**).

Taken together, these results support that the ABE and CBE are useful in creating specific point mutations in the homologous *HBG1* and *HBG2* promoters, leading to a significant increase in the number of HbF positive cells and overall HbF production.

## DISCUSSION

During normal globin switching, interactions of cis-acting elements with several different transcription factors lead to the silencing of fetal globin and, in turn the activation of beta-globin (23). To obtain insights into the regulation of gamma-globin gene expression, we have used two complementary base editing approaches to screen the *HBG* promoter at single-nucleotide resolution. This approach allowed us to identify several novel nucleotide substitutions in the *HBG* promoter that elevate HbF levels.

Current approaches to study fetal globin regulation by programmable nucleases often result in the deletion of the *HBG2* gene due to the introduction of DSBs in both *HBG* promoters (24). The elimination of the 4.9 kb intergenic region (including *HBG2*) favors the locus control region (LCR) to directly interact with *HBG1* promoter and regulate its expression without competing with the *HBG2* promoter (12). Hence, it is challenging to determine the exact role of different HPFH mutations on individual gamma-globin expression because mutations can occur in both *HBG2* and *HBG1* promoters. On the other hand, a base editing strategy converts bases at the target site without the generation of DSBs and hence avoids splicing of the *HBG* locus. Using this strategy, we targeted regions in both *HBG1 and HBG2* promoters and were able to efficiently edit sites in the promoters without causing deletion, which gave us the opportunity to evaluate gamma-globin expression from two active promoters.

Several different HPFH point mutation has been reported in *HBG* promoter; however, the effect of these mutations on gamma-globin expression in the native cellular environment has been deciphered only for few mutations (1,7,8,15,19,20). In this study, we have identified several new point mutations in the *HBG* promoter associated with high HbF levels. Our findings are in agreement with previous reports that the point mutations in three different regions of the *HBG* promoters centered around positions -198, −175, and −115 mimic the non-deletional HPFH point mutations which are essential regulators of HbF expression (5,7,8,15,19,20). In addition to the known mutations, we have identified novel substitutions at -69 (C to T), -70 (C to T), -122 (G to A), -123 (T to C) and -124 (T to C) of the *HBG* promoters as potential new regulatory mutations that can elevate gamma-globin expression. The levels of gamma-globin expression resulting from these mutations were very similar to that of well-characterized HPFH mutations. Further work needs to be done to identify the transcriptional regulators that are responsible for gamma-globin silencing or activation in these regions.

*HBG* promoter base editing by ABE mediated conversion (A to G) revealed multiple potential HbF regulatory regions compared to CBE since the targeted region had more ABE compatible gRNAs than CBE. Base conversion within the -115 cluster (from -110 to -116) showed the highest increase in promoter activity, confirming previous studies (25–28). CBE mediated base conversion (C to T) at position -114 and -115 resulted in significant induction of HbF than the multiple A to G nucleotide substitutions at -110, -112, -113, -116 positions made by ABE. Recently, it has been shown that the major HbF repressor BCL11A directly binds to the core TGACC motif located at – 114 to -118 (19). Naturally occurring HPFH mutations at -117G>A, -114C>A, -114C>T, -114C>G, and a 13bp deletion (Δ13bp) disrupt binding of BCL11A to the promoter (8). The -113A>G HPFH mutation within the -115 cluster creates a binding site for the master erythroid regulator GATA1 without disrupting the binding of BCL11A (7). Our results are consistent with these previous reports showing that disruption of the core binding region of BCL11A and the creation of a *de novo* binding sites for GATA1 results in the elevation of fetal globin in wild type HUDEP-2 cells (7). ABE mediated T to C substitution at position -198 of the *HBG* gene promoter has previously been shown to be associated with British HPFH and substantially elevate expression of HbF by creating a *de novo* binding site for the erythroid gene activator Kruppel-like factor 1 (20,29). Another known HPFH mutation (−175T>C) has been shown to promote enhancer looping to the *HBG* promoter through recruitment of the activator TAL1 (15). Besides, -173 and -175 mutations disrupt the binding of GATA1 and OCT1, respectively, and reactivate *HBG* expression (30). Interestingly, editing of the -175 T>C position by ABE with a single gRNA, resulted in a dramatic increase in HBG expression even with relatively low editing efficiency (20 %). Prior findings demonstrated that the nuclear factors bind to four repeating copies of CCTTG present in the region of the HBG promoter (from -97 to -126 of TSS) and limits the binding of KLF1, leading to suppression of HBG expression(31). In this study, we identified novel single nucleotide substitutions (−122 (G to A), -123 (T to C), -124 (T to C)) at one of the CCTTG region (−122 to -126) in the *HBG* promoter that reactivates gamma-globin expression. We anticipate that these induced novel mutations may allow KLF-1 binding in the target region and reactivate gamma-globin expression, as demonstrated by the previous study. Future work will focus on unraveling the molecular mechanisms that are responsible for this reactivation.

DNA methylation is a dynamic epigenetic mark mostly found on cytosine residues of certain CpG dinucleotides in mammalian genomes (32). The overall methylation patterns of the *HBG* promoter regions are inversely related to gene expression (33). In six CpG sites flanking the transcription start site of *HBG*, the methylation status of the −53 CpG has been shown to correlate with gene expression in erythroid cells (33). However, CBE mediated conversion of C to T in the -53 region of the *HBG* promoter did not show any considerable effect on globin expression, suggesting that methylation of this CpG dinucleotide is not critical for sustained HbF repression.

The translational potential of genome-edited hematopoietic stem and progenitor cells depends on long-term engraftment and repopulation ability. However, genotoxicity and cytotoxicity that can arise as a result of DSBs generated by programmable nucleases can be a huge limiting factors (34,35). A previous study in nonhuman primate observed that the *HBG* promoter editing by Cas9 resulted in *HBG2* deletion with up to 27% frequency and that cells with this deletion had a selective disadvantage during engraftment (36). Base editing at the target sites of *HBG1* and *HBG2* promoter by ABE and CBE does not result in any large deletions of the intergenic region and only showed very indel formation. ABEs have an inherent advantage over CBEs as it generates desired edits (A:T to G:C) only, whereas the latter generate unanticipated edits. In corroboration with existing findings, our results also suggest that ABE is the better base editor than CBE with respect to purity of base conversion and indel formation (37). Our proof of principle study validated various gRNAs that can elevate the HbF levels to therapeutic levels laying the groundwork for potential clinical applications. This approach could address a range of ß-globin disorders avoiding the need to develop specific therapeutic products for each of them.

## CONCLUSION

In summary, we have demonstrated that CRISPR base editing can be utilized to drive the expression of fetal hemoglobin to therapeutically relevant levels in an erythroid progenitor cell line. After screening every gRNAs within the 320 bp region of the *HBG* promoter, we identified nine gRNAs that, when paired with the appropriate base editor, can introduce HPFH-like mutations without the generation of DSBs. We identified five novel regulatory regions for HBG1/2 that are involved in the silencing of gamma-globin in adult erythroid cells shedding light on the molecular mechanisms behind hemoglobin switching (**Fig. 6**). Furthermore, we think that our base editing strategy may lead the way to novel therapeutic strategies for the β-hemoglobinopathies in the future.

**Figure 6:**
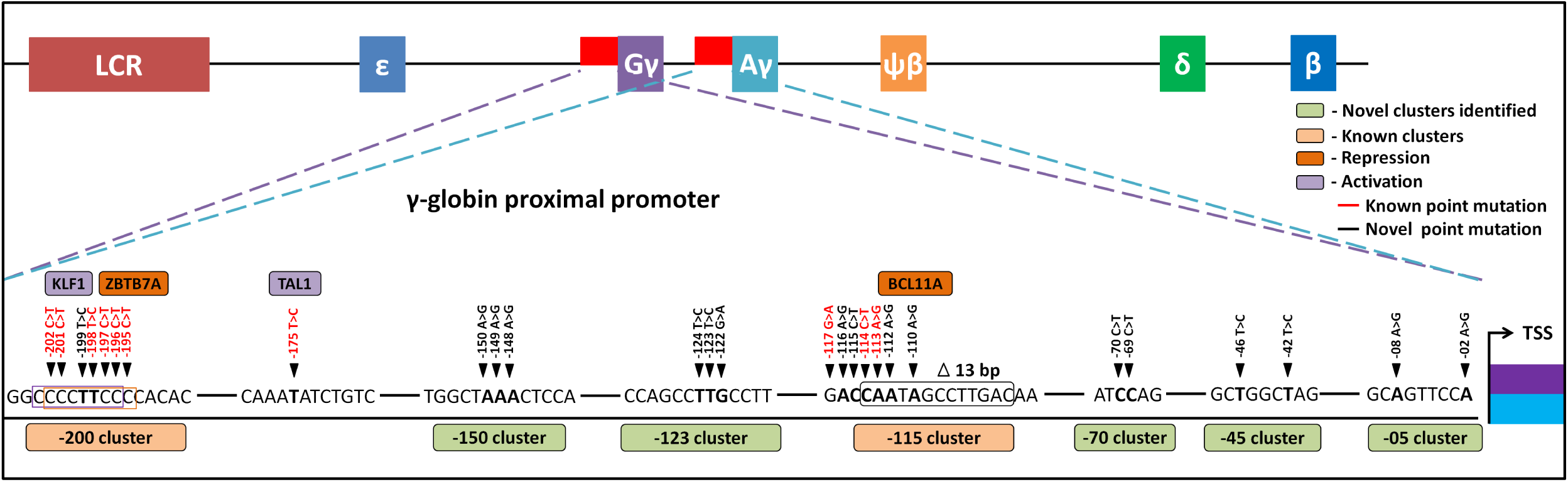
Schematic representation of novel target sites identified in the HBG1/G2 promoter. The proximal promoter region of HBG2 and HBG1 are represented from TSS till -205 bases. Novel clusters identified from this study are highlighted in Sage (5 clusters), and known clusters are highlighted in Melon - (2 clusters). In those clusters, the black triangular notches represent single base conversion responsible for novel HPFH like point mutations (in black font) and well-known HPFH point mutations (red font). Among all the represented point mutations, the novel base conversions from our study are represented in bolded font, while the known bases from other studies are represented in normal fonts. Transcriptional activators (Lavender) and repressors (Orange) that bind to the known clusters are also depicted in the figure.

## MATERIALS AND METHODS

### Designing and cloning of the gRNA

The guide RNAs (gRNAs) for targeting the *HBG1/G2* promoter region spanning from -1bp to -320 bp were designed using the SnapGene and Benchling software. The criteria for choosing the gRNAs and cloning were mentioned in supplemental methods. The synthetic complementary oligonucleotides listed in **Sup.table.1** were annealed (38,39) and cloned into *BsmBI* digested pLKO5.sgRNA.EFS.GFP vector (40)gift from Benjamin Ebert ((Addgene #57822)). The ligated product was transformed into DH10B competent cells (41). Positive clones were confirmed by colony PCR and then validated by Sanger sequencing.

### Plasmid Constructs

The plasmid in this study, pLenti-FNLS-P2A-Puro (Addgene#110841-CBE) and pLenti-ABERA-P2A-Puro (Addgene#112675-ABE) was a gift from Lukas Dow (42), pMD2.G and psPAX2 (Addgene #12259, 11260-2nd generation lentiviral packaging construct)was a gift from Didier Trono. The plasmids were isolated using NucleoBond Xtra Midi EF (Macherey-Nagel) according to the manufacturer’s instruction.

### Lentivirus production

About 1×10^6^ Lenti-X™ 293T cells were seeded in a 10cm cell culture dish with DMEM (HyClone) containing 10% FBS and 1 % Pen-Strep. After the 80% confluency, 2.5 μg of pMD2.G, 3.5 μg of psPAX2 along with 4 μg of lentiviral-vector (base editing or sgRNA constructs) were transfected using FuGENE-HD as per manufacturer’s protocol. Viral supernatant was then collected and concentrated using Lenti-X Concentrator (Takara). The concentrated pellet was resuspended in 200 μl 1xPBS.

### HUDEP-2 transduction and erythroid differentiation

HUDEP-2 cell lines were cultured in StemSpan™ SFEM II (STEMCELL Technologies) along with 50 ng/ml SCF (ImmunoTools), 3 U/ml EPO (Zyrop 4000 IU Injection), 1x Pen-Strep, 1 uM dexamethasone (Alfa Aesar), 1 μg/ml doxycycline (Sigma-Aldrich) and 1x L-Glutamine 200 mM (Gibco™) (16). The lentivirus containing the pLenti-FNLS-P2A-Puro or pLenti-ABERA-P2A-Puro or desired gRNAs were added to the 0.5 million of HUDEP-2 or HUDEP-2 stable cell line in a one well of the six-well plate along with 6 μg/ml Polybrene (Sigma-Aldrich) and 1% HEPES 1M buffer (Gibco) and centrifuged at 800g for 30 min at RT. In case of gRNA transduction, the transduction efficiency was analyzed by GFP expression using FACS. For the stable cell line generation, the cells were then treated with 1 μg/ml puromycin (Sigma-Aldrich) after 48 hours for ten days. For the erythroid differentiation, we followed the protocol as mentioned in the supplemental method (43).

### Base editing efficiencies and Sequencing

To determine the base editing efficiency, the genomic DNA from the edited cells was isolated after the transduction for eight days using a DNA isolation kit (NucleoSpinBlood - Macherey-Nagel). The target region for base editing was amplified using the respective primers **(Sup Tables 2 and 3)**, and the amplified product was Deep, or Sanger sequenced and analyzed as mentioned in supplemental methods.

### Real-time PCR

Total mRNA of edited HUDEP-2 cells was isolated using the NucleoSpin RNA kit (Macherey-Nagel) and reverse transcribed into cDNA. Quantitative real-time PCR (qRT-PCR) was carried out using respective primers **(Sup. Table 2)**, and the expression levels *of HBB, HBA*, and *HBG* mRNA were determined as mentioned in supplemental methods.

### HbF intracellular staining

To evaluate the frequency of HbF positive cells, the expanded or differentiated HUDEP-2 cells were fixed, permeabilized, and HbF intracellular staining was performed (44), as mentioned in supplemental methods. Finally, the stained cells were analyzed for HbF positive cells using FACS.

### Hemoglobin HPLC

The differentiated cells were collected and washed with 1x PBS and resuspended in 1100 μl cold ddH20. The cells were sonicated for 30 seconds with 50% Amp in ice and centrifuged at 14000 rpm for 15min at 4°C. The supernatant was analyzed for hemoglobin in the VARIANT II Hemoglobin Testing System (Bio-Rad). Reverse-phase HPLC was performed on Shimadzu Corporation-Phenomenex for globin chain quantification(45).

### Statistics

The statistical tests were performed using GraphPad Prism 8.1 and Microsoft Excel. Since all the data were normally distributed, unpaired two-sided t-test or one-way ANOVA was carried out as appropriate. In all the tests, p<0.05 was considered statistically significant. Linear regression was carried out to find out if any correlation exists between two variables.

## ABBREVATIONS

ABE: Adenine base editor
CBE: Cytosine base editor
HUDEP-2: Human umbilical cord blood- derived erythroid progenitor-2
HPFH: Hereditary persistence of fetal hemoglobin
HbF: Fetal hemoglobin
HBG: Gamma-globin
TSS: Transcription start site
HDR: Homology-direct repair
Indels: Insertion and deletions
NHEJ: Non-homologous end joining
DSBs: Double-strand DNA breaks

## ACKNOWLEDGMENTS

The research reported in this work was supported by NAHD grant: BT/PR17316/MED/31/326/2015 (Department of Biotechnology, New Delhi, India), EMR grant: EMR/2017/004363 (Science and Engineering Research Board (SERB), New Delhi, India), and Indo-U.S. GETin Fellowship_2018_066 (Indo-U.S. Science & Technology Forum (IUSSTF)). We sincerely acknowledge CSCR (a unit of inStem, CMC Campus, Vellore, India) for providing the startup funds. B.W. is supported by an Early Career Research Fellowship from the National Health and Medical Research Council Australia. We thank Bandlamudi Bhanu Prasad, Anila George, Nazar Syed Basha, Nivedhitha Devaraju, Vignesh Rajendran, Kirti Prasad, and Dr. Muthuraman. N, CSCR, Vellore, for the technical assistance and helpful discussions. We thank Mrs. Sumithra and Mr. Neelagandan at the Department of Hematology, CMC, for help with HPLC variants and Keerthivasan.R.C, IISER Mohali, for bioinformatics. Visual abstract was made using BioRender.com. Also, we like to acknowledge the CSCR core facility for supporting us with all the required instrumentations.

## AUTHOR CONTRIBUTION

M.K.M. conceived the project and designed the experiments with support from J.E.C., B.W., M.A.D., and N.S.R; N.S.R performed the majority of the experimental work with advice from M.K.M., B.W., and M.A.D; N.S.R., and M.K.M. analyzed the data; Y.N., and R.K. provided HUDEP-2 cells; S. K. W., and J.V. performed NGS analysis; A.A., and B.P., performed HPLC chain analysis; J.E.C., B.W., M.A.D., A.S., R.V.S., S.T., and S.M., provided intellectual insight and feedback on the data and manuscript; N.S.R., and M.K.M. wrote the manuscript, and all authors read and approved the contents of the manuscript.

