## Supplementary information for "Identification of novel HPFH-like mutations by CRISPR base editing that elevates the expression of fetal hemoglobin"

### TABLE OF CONTENTS:


1. Visual Abstract
2. Supplemental methods
   i. Guide RNA designing
   ii. Cloning
   iii. Editing efficiency and sequencing
   iv. HbF intracellular staining
   v. Real Time PCR
   vi. Erythroid differentiation
   vii. High-performance liquid chromatography
3. Supplemental Reference
4. Supplemental Figures
   – Figures 1 to 9
5. Supplemental legends
6. Supplemental Tables
   – Tables 1 to 3

#### VISUAL ABSTRACT

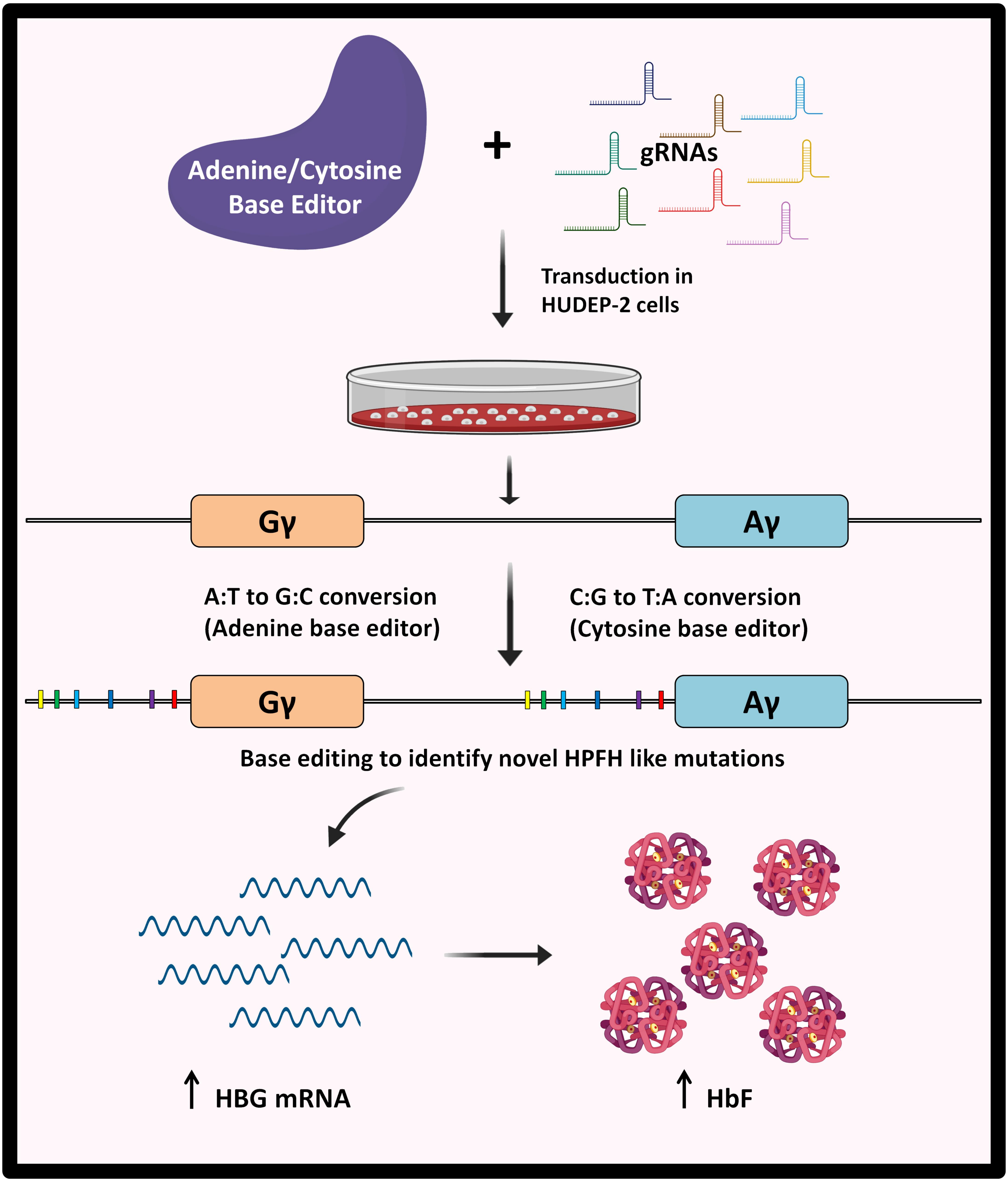

### SUPPLEMENTAL METHODS

#### Guide RNA designing:

First, we designed gRNAs for CBE using design type gRNAs for "base editing" in the Benchling tool and got around 43 hits from which we selected 32 non-overlapping gRNAs. Since there was no design type gRNA for ABE in the Benchling tool, we manually designed gRNAs using SnapGene software. Among the 32 gRNAs selected for CBE, 27 gRNAs also had an editing window compatible with ABE. We manually designed another nine gRNAs for ABE using SnapGene software, apart from the gRNAs that were obtained from the Benchling tool. The forward primer consists of the gRNA sequence without PAM (20 bp) and "CACCG" overhang at the 5' end, while the reverse primer consists of "AAAC" overhang at the 5' end, a reverse complement of gRNA without PAM (20 bp) and a "C" added at 3' end.

#### Cloning:

The vector backbone was prepared by digesting pLKO5.sgRNA.EFS.GFP plasmid using the *BsmBI* enzyme (NEB) according to the manufacturer's instruction and then gel purified using Zymoclean Gel DNA Recovery Kit. The oligo-annealed products were diluted 1:200 fold, from which 6 µl was taken along with 50 ng of vector backbone and ligation reaction was set up as per the manufacturer's instruction from NEB. The ligated product was transformed into DH10B competent cells and plated in LB-agar containing 100 µg/ml of Ampicillin for selection.  Three colonies were picked from the plate and inoculated in LB broth for colony PCR.  Colony PCR was carried out using GoTaq® Hot Start Polymerase premix (Promega) and 1 µl each of forward and reverse sequencing primers (10 picomoles) along with 1 µl of processed cells (Sup. Table 2) in a thermocycler (Applied Biosystems® Veriti®). The cyclic conditions were as follows: initial denaturation at 95°C for 10 min, 35 cycles of 95°C for 30 secs, 55°C for 30 secs, 72°C for 45 secs, followed by a final extension at 72°C for 7 min. After confirming the expected amplification in 1% agarose gel, Sanger sequencing was carried out using 10 µl of the amplified product.

**Editing efficiencies and Sequencing:**

The genomic DNA was isolated from the edited samples using a DNA isolation kit (NucleoSpin Blood - Macherey-Nagel) and used to set a PCR using GoTaq® Hot Start Polymerase premix, which amplifies the promoter regions of *HBG1* and *HBG2*. The Primers used were HBF 1 forward, HBF 2 forward, and HBF 1 Reverse primer, as mentioned in Sup. Tables 2 and 3. The cyclic conditions were as follows: initial denaturation at 95°C for 10 min, 35 cycles of 95 °C for 30 secs, 60 °C for 30 secs, 72 °C for 45 secs, followed by final extension of 72 °C for 7 min. Once the amplification was confirmed using agarose gel electrophoresis, it was subjected to Sanger sequencing. Amplicon sequencing was performed using the MiSeq System (Illumina) for the PCR product obtained by amplifying the promoter regions of *HBG* by GXL premix as per manufacturer's instruction. The Primers used were NSG 2, NSG 3, and NGS 4, as mentioned in Sup. Tables 2 and 3. The protocol for library preparation and sequencing was from the Corn Lab, Innovative Genomics Institute (IGI), UC Berkeley (1). The Fastq files obtained were analyzed for base editing using CRISPResso-2 (2). NGS 4F and NGS 2R primers were used to to differentiate editing efficiency between *HBG1* and *HBG2* for triplicate samples using Bowtie2 and IGV (3,4). Incase of Sanger sequencing data, tools like CRISPR Edits (ICE) (Synthego) and EditR were used to find out the indels and the base editing efficiency, respectively (5,6).

**HbF intracellular staining:**

Edited cells were spun at 1000 rpm for 5 min at RT. The pellet was resuspended in 200µl BPBS (containing 1xPBS and 0.1% BSA) and transferred to a 96 well cell culture plate. The plate was spun at 1000rpm for 5min, and then the pellets were resuspended in freshly prepared 100µl of 0.05% glutaraldehyde (MP Biomedicals) using BPBS and incubated at RT for 8 min. Each well was topped up with 100 µl of BPBS, spun at 1000 rpm for 5 min, and the pellet was washed with 200µl BPBS. 100µl of 0.1% fresh Triton x-100 (Fisher scientific) prepared in BPBS was then added to the pellet, resuspended, and incubated at RT for 7 min. After 7 minutes, 100µl of BPBS was added, centrifuged at 1000 rpm for 5min, and the pellet was again washed with 200 µl BPBS. 50µl of BPBS containing 2µl of HbF antibody (Fetal Hemoglobin Monoclonal Antibody (HbF-1), APC - Thermo Fisher) was added to each well except the unstained and kept in the dark for 15min. After topping it up with 100µl of BPBS, the plate was spun at 1000 rpm for 5min, the supernatant was discarded, and the pellet was washed again with 200µl BPBS. The pellet was resuspended in 300 µl of BPBS and taken for FACS analysis (BD FACS ARIA-III) to check for HbF positive cells based on APC fluorochrome staining.

**Real-time PCR:**

RNA isolation was performed with the NucleoSpin RNA kit (Macherey-Nagel) and quantified using Nanodrop (Thermo Fisher Scientific), from which 1 µg was taken for cDNA conversion. Reverse transcription was performed using the iScript™ cDNA Synthesis Kit (Bio-Rad) as per the manufacturer's protocol in a Veriti™ PCR system. A negative RT reaction was also set up to check for genomic DNA contamination. The expression levels of *HBB*, *HBA*, and *HBG* (Sup. Table 2) were determined by qRT-PCR using SsoFast™ EvaGreen® Supermixes (Bio-Rad) and a QuantStudio™ 6 Flex Real-Time PCR System (Applied Biosystems). The qRT-PCR mixture (10 µl) contains 1µl each of respective forward and reverse primer (5 µM), 5 µl of SYBR green master mix, 2µl of H20, and 1µl of 5 fold diluted cDNA template. *GAPDH* (Sup. Table 2) was used as an internal control gene to normalize the data for ΔΔCT (relative expression analysis). The cycling condition was performed as per the manufacturer's protocol (Bio-Rad). A dissociation curve analysis was carried out to ensure there is no unspecific amplification.

**Erythroid Differentiation:**

Around 1 million transduced cells were taken and used to setup differentiation after eight days of expansion. The edited cells were seeded in a 65mm dish containing 5 ml of differentiation media and were differentiated for nine days with media change on day 3 (cells are taken to 10 cm dish) and day 6. The media consists of IMDM glutamax (Gibco®), 3% AB serum (MP Biomedicals), 2% FBS, 0.1% Insulin solution human(Sigma-Aldrich), 3 U/ml Heparin sodium salt (MP Biomedicals), 200 µg/ml Holo Transferrin (BBI Solutions), 3 U/ml EPO, 10 µg/ml SCF, 1 ng/ml IL3 (Immuno Tools), 1x Pen-Strep and 1 µg/ml Doxycycline. Day 6 media was prepared devoid of Doxycycline, and it consists of 500 µg/ml of holo transferrin instead of 200 µg/ml. After nine days of differentiating the cells, flow cytometry analysis of CD235 and CD71 markers was performed to find the differentiation pattern.

**High-Performance Liquid Chromatography (HPLC):**

The transduced cells after nine days of differentiation were collected and washed with 1x PBS and resuspended in 1100 µl cold nuclease-free water. The cells were continuously sonicated using a probe sonicator for 30 seconds with 50% Amp in ice. The sonicated product was centrifuged at 14000 rpm for 15 min at 4$^{o}$C. After centrifugation, 1000 µl of supernatant was stored in -80 $^{o}$C for the VARIANT II Hemoglobin Testing System (Bio-Rad), and the other 100 µl was also stored in -80 $^{o}$C for Reverse phase HPLC (Shimadzu Corporation-Phenomenex) for globin chain quantification. Globin chains were measured using a Shimadzu UFLC consisting of binary gradient pumps (LC-20A), autosampler (SIL-HTC), and a column oven (CTO-20AC) coupled with UV detection (Shimadzu™, Kyoto, Japan). The data was analyzed using LC Solutions™ software. Chromatographic separation of the analytes was done using Aeris Widepore 3.6 lm XB-C18 25cm 4.6mm column behind a SecurityGuard UHPLC Widepore C18 4.6mm guard column (Phenomenex™). The LC conditions were as follows: Solvent A: 0.1% Trifluoroacetic acid (TFA), pH 3.0 and Solvent B: 0.1% TFA in acetonitrile was used as mobile phase with gradient elution (40% Solvent B) at a flow rate of 1.0ml/min and column temperature maintained at 70 C. The total run time is 8 minutes. The UV detection was set at 190nm for globin chain detection.

### SUPPLEMENTAL FIGURES

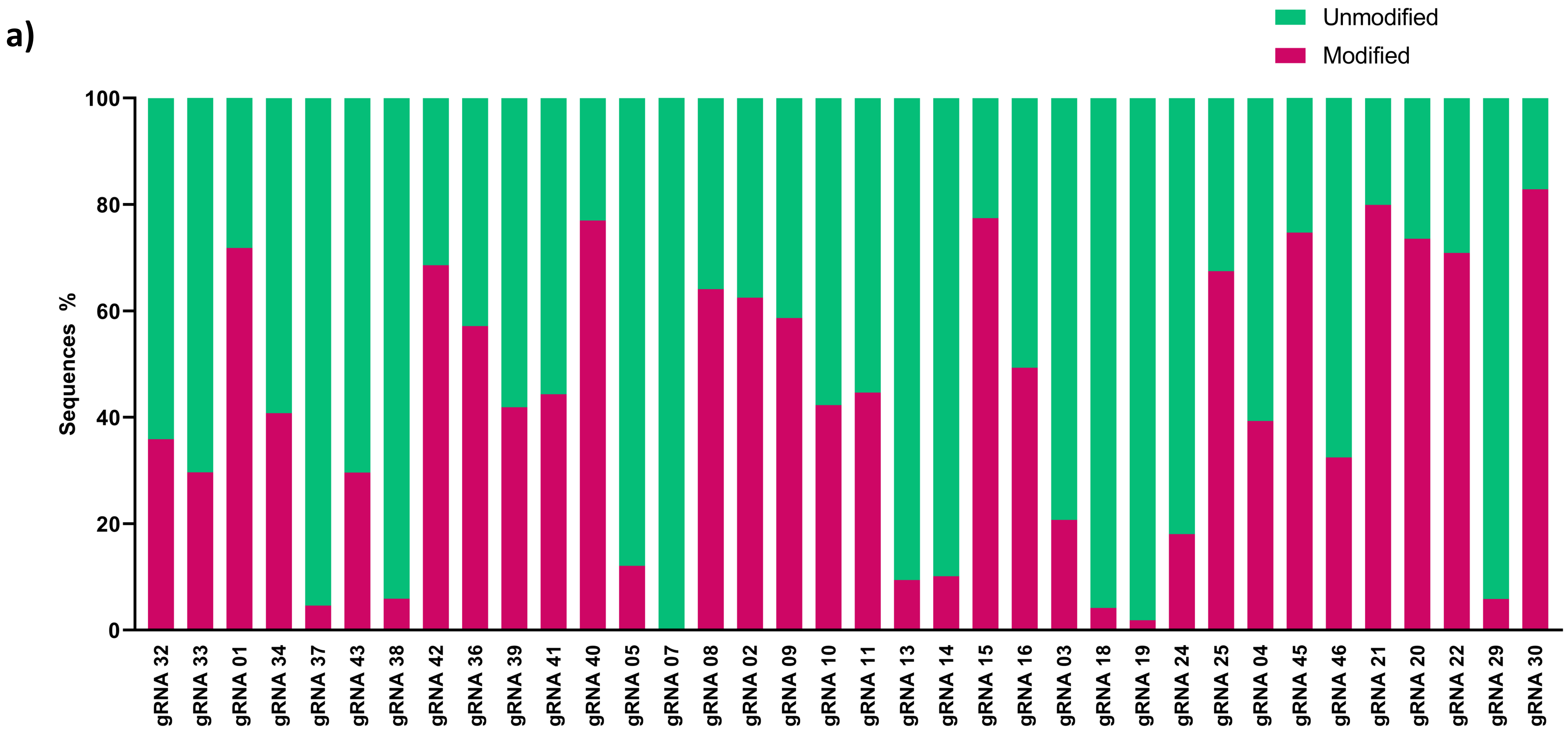


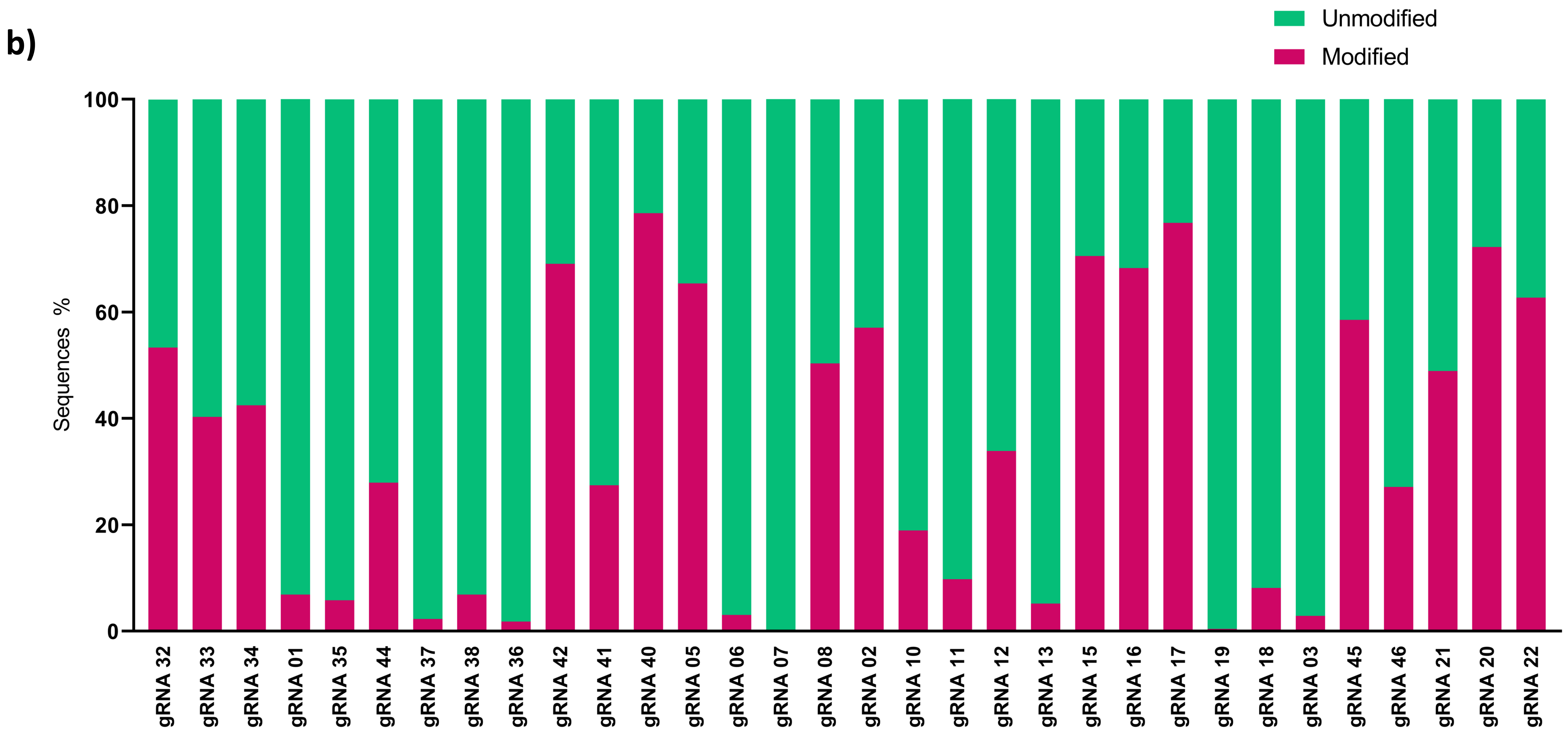


#### Supplementary Figure 1

a)

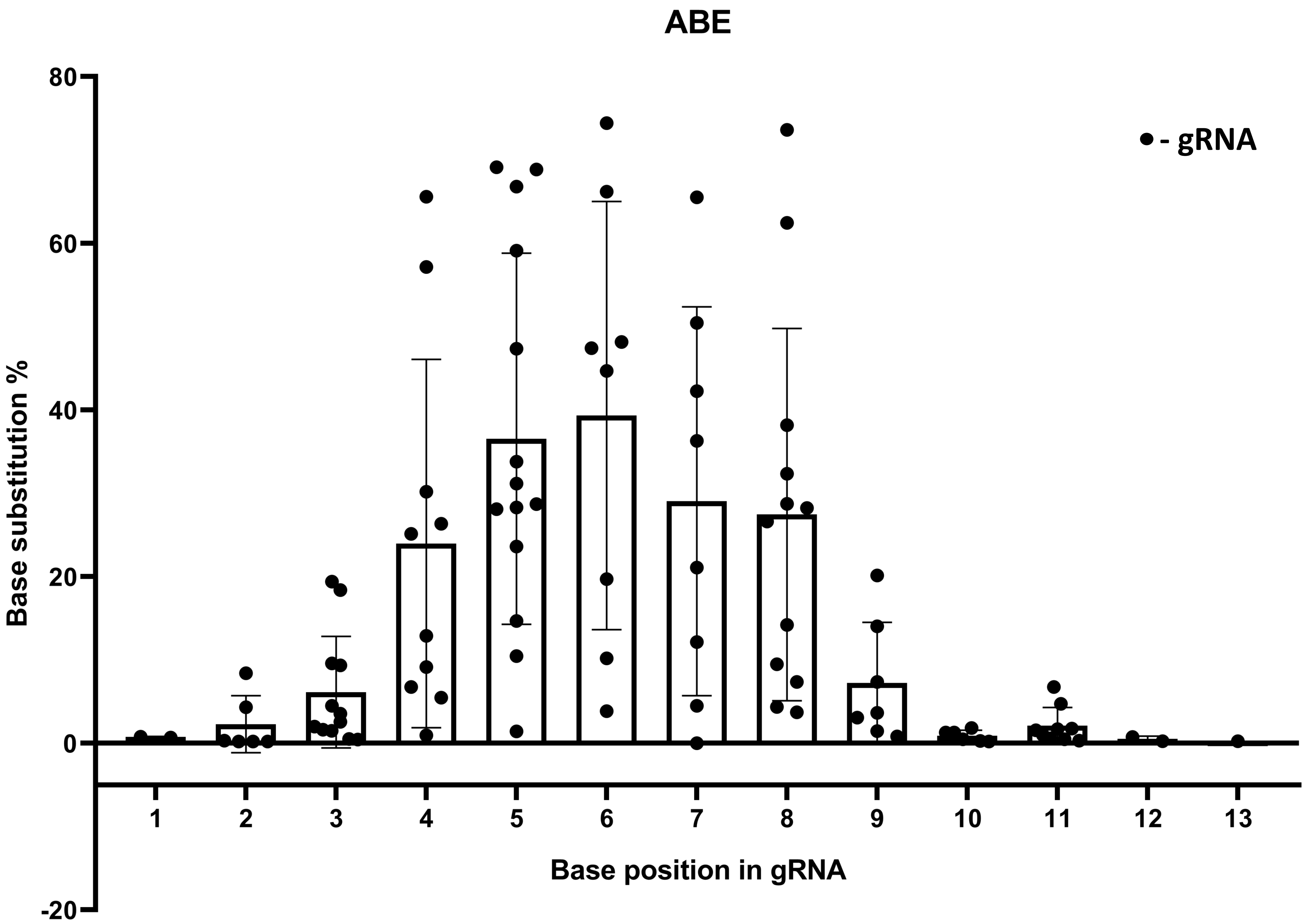


b)

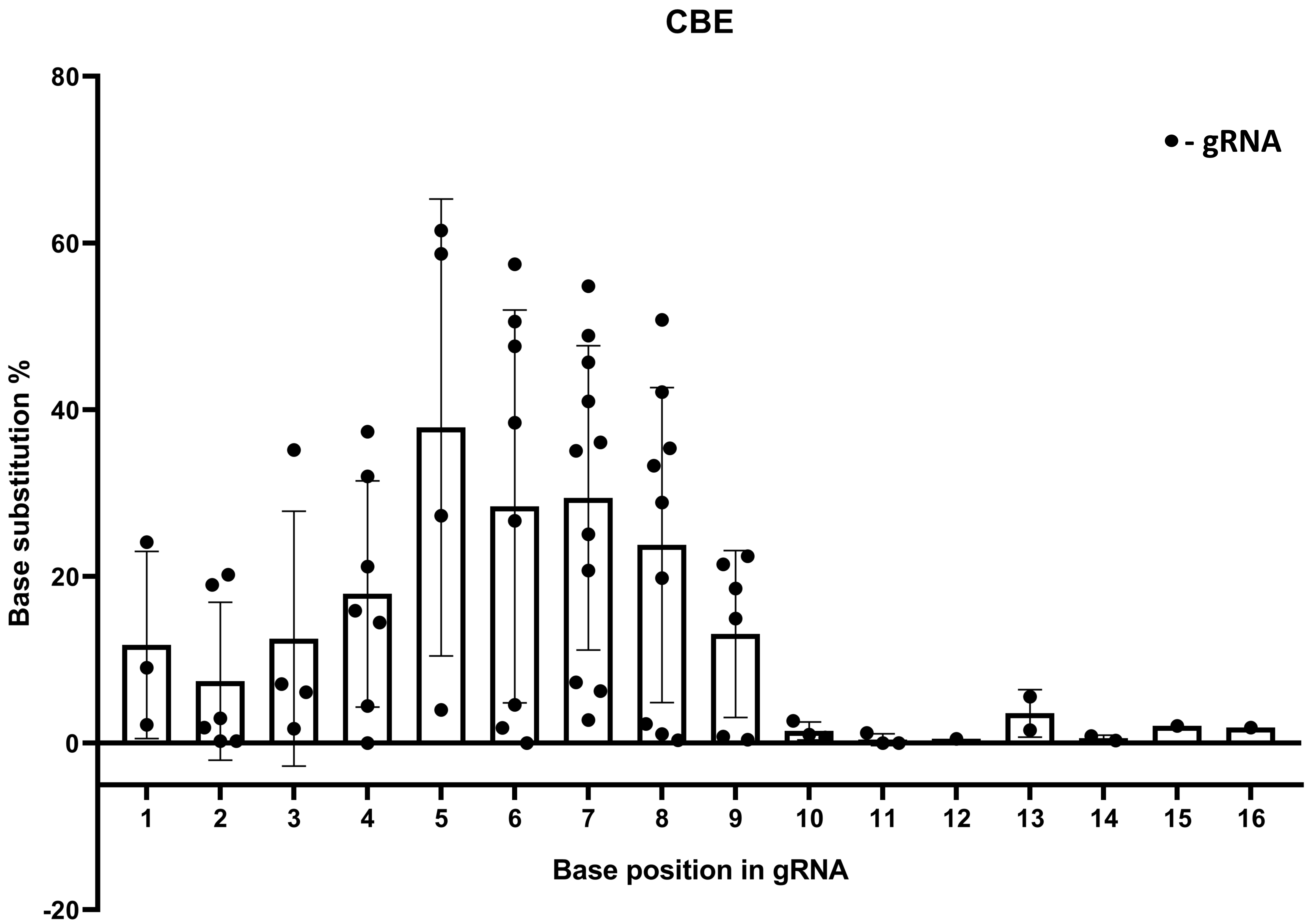


**Supplementary Figure 2**

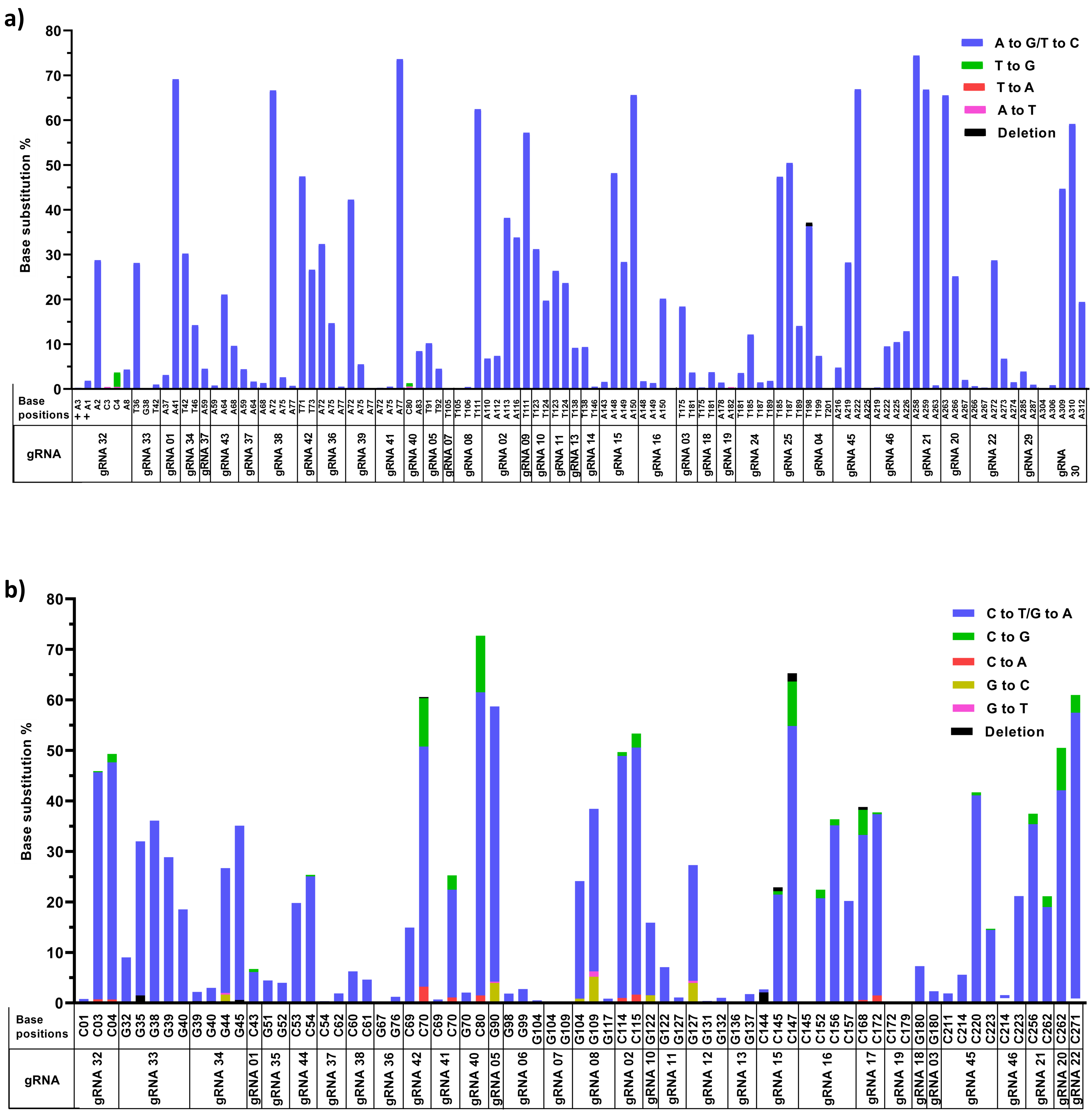


#### Supplementary Figure 3

a)

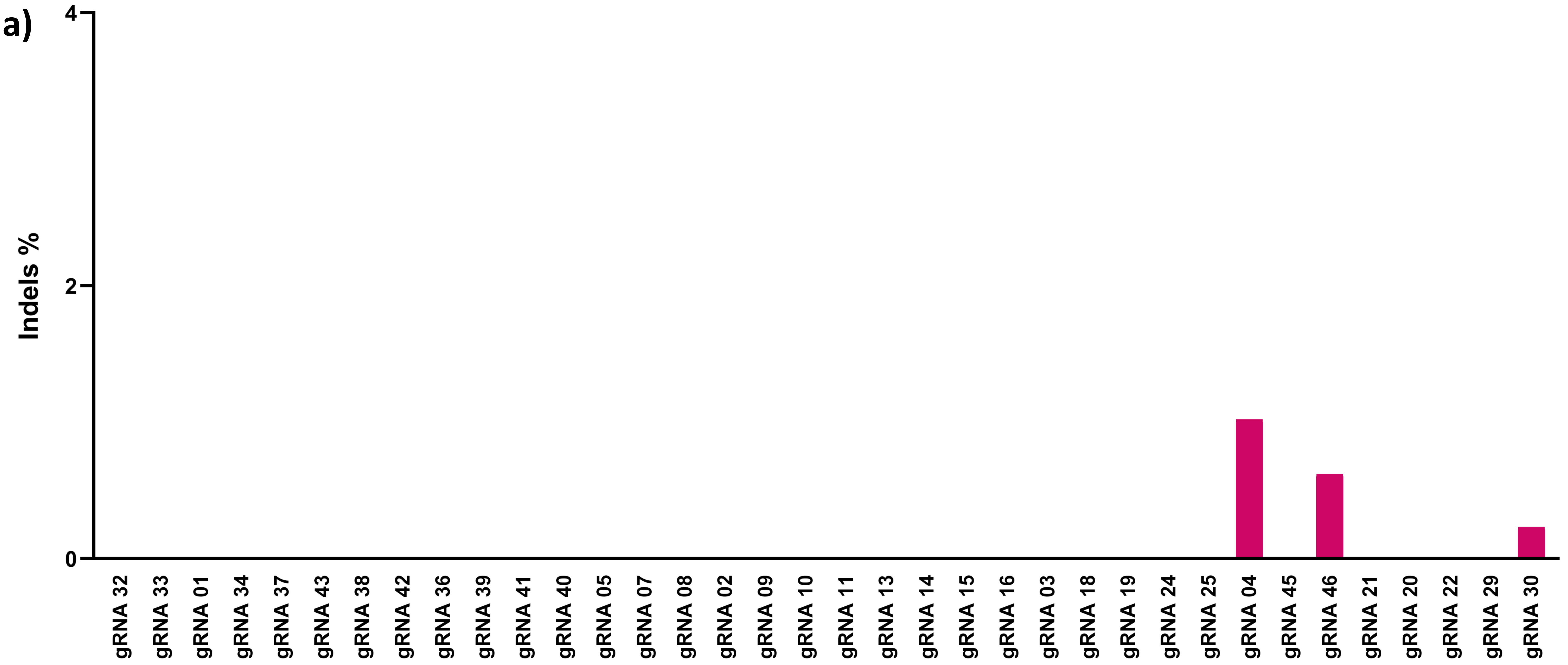


b)

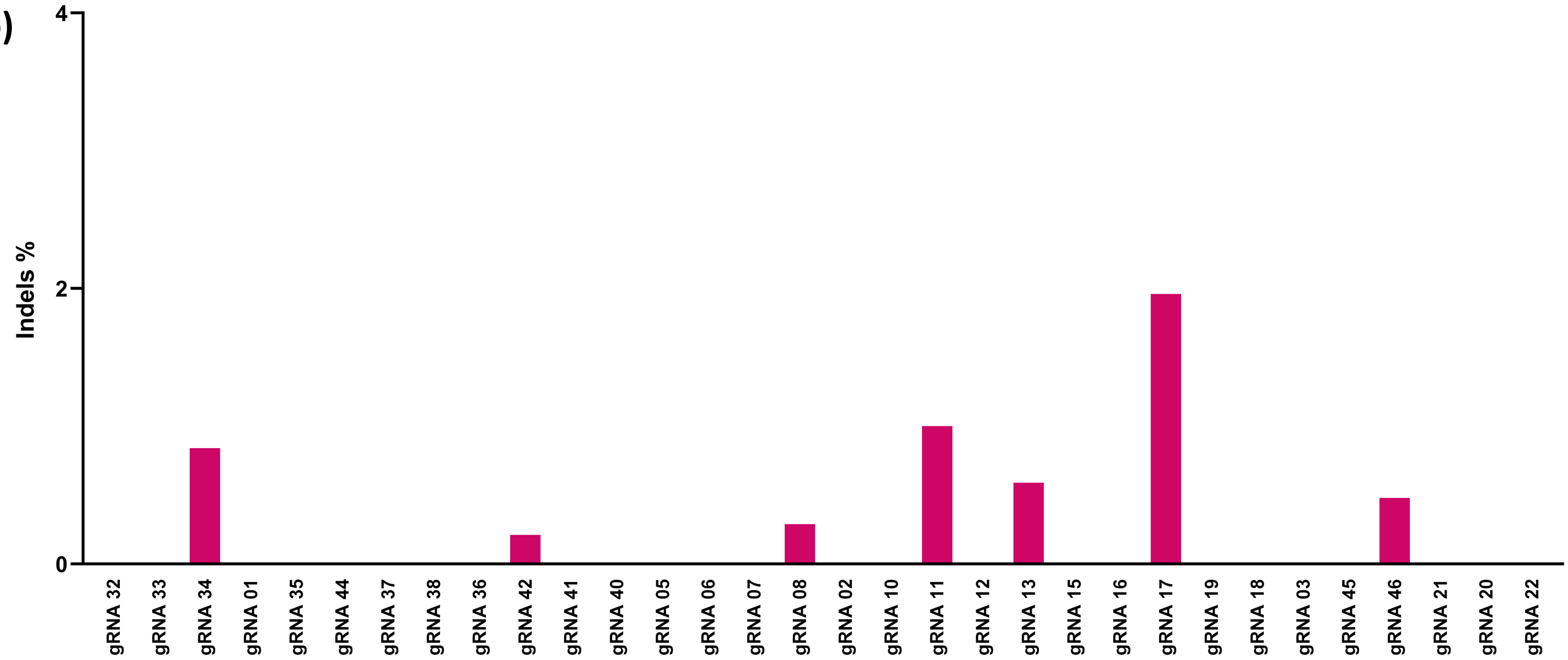


#### Supplementary Figure 4

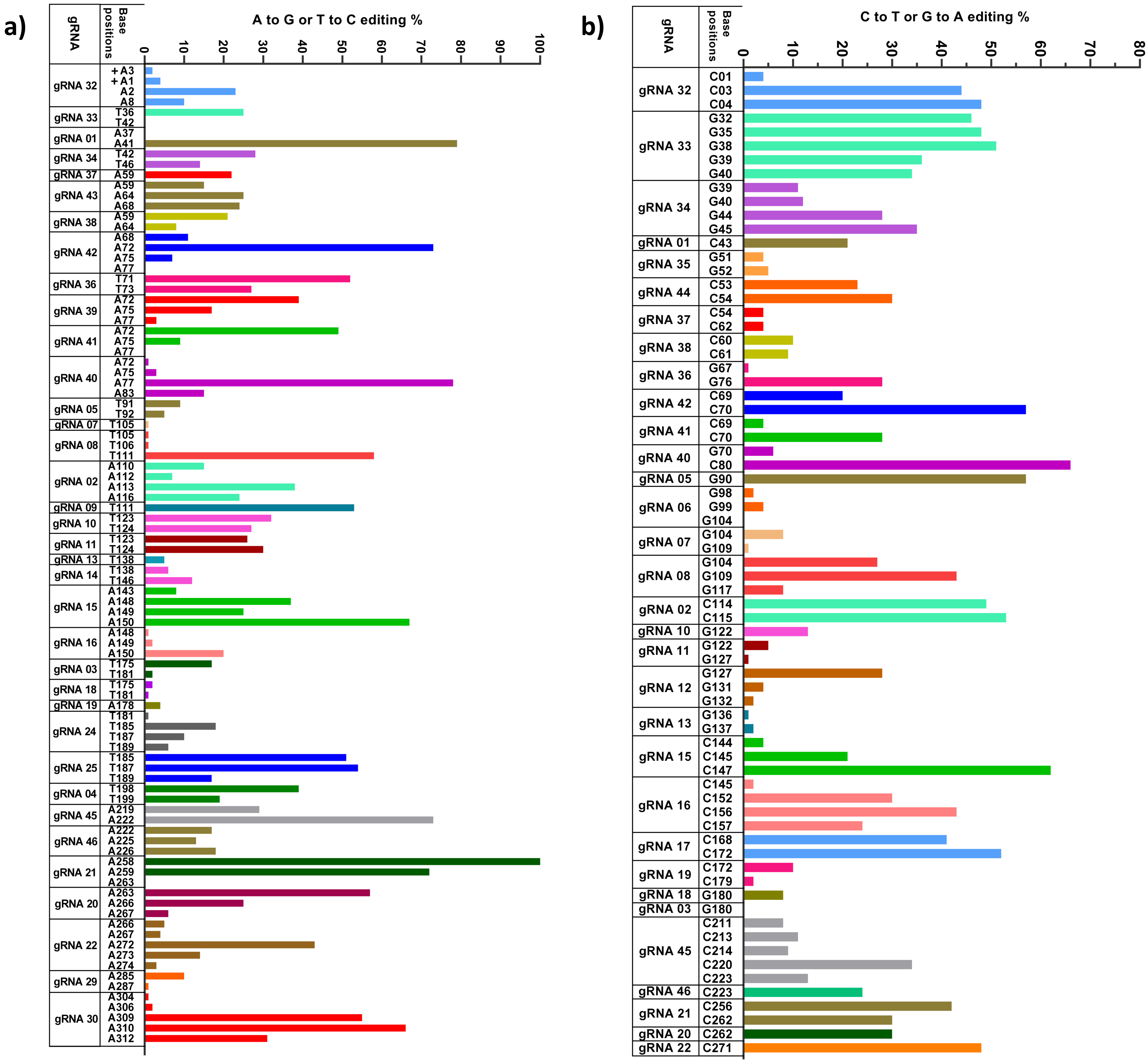


#### Supplementary Figure 5

a)
b)
c)
d)
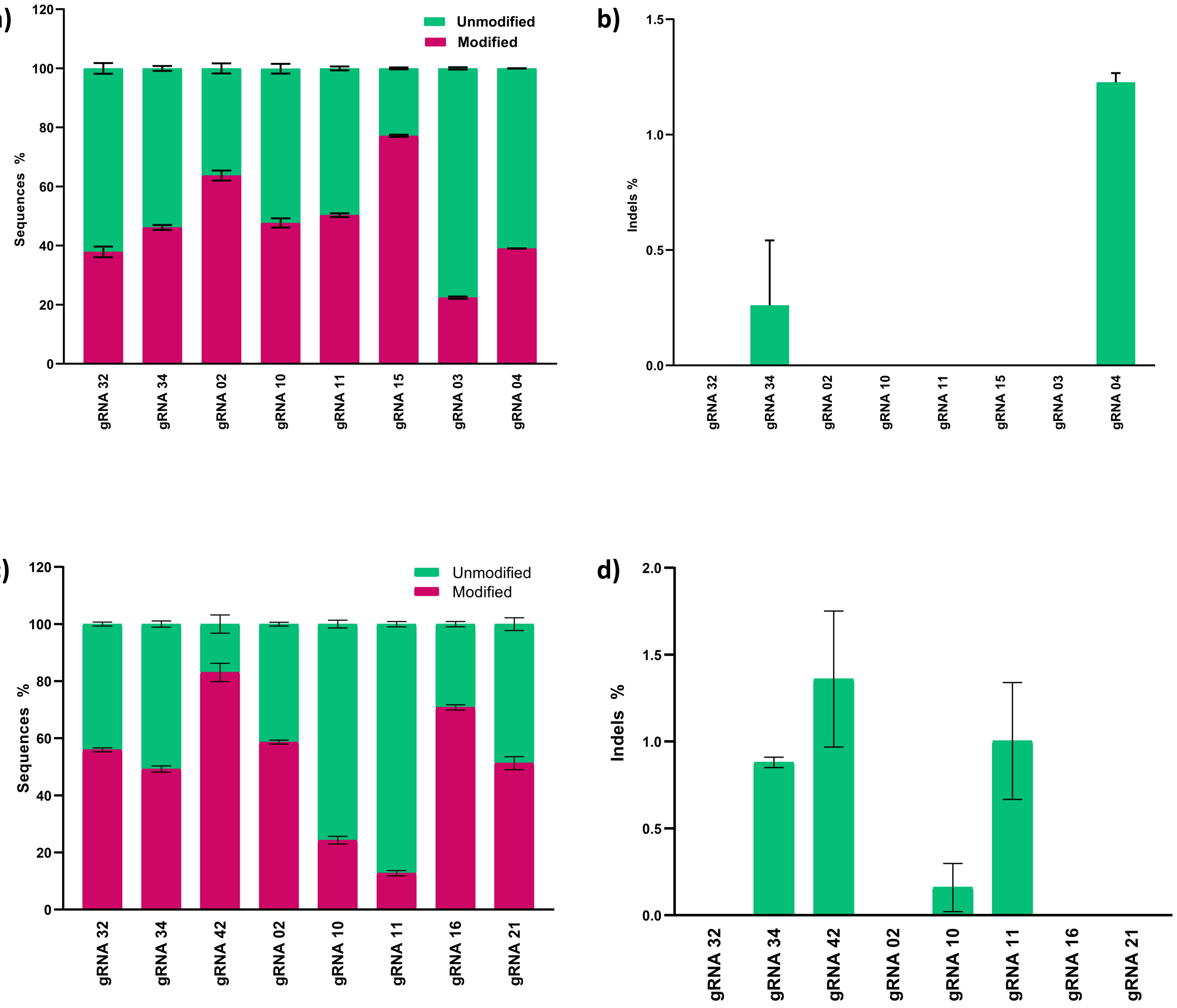


#### Supplementary Figure 6

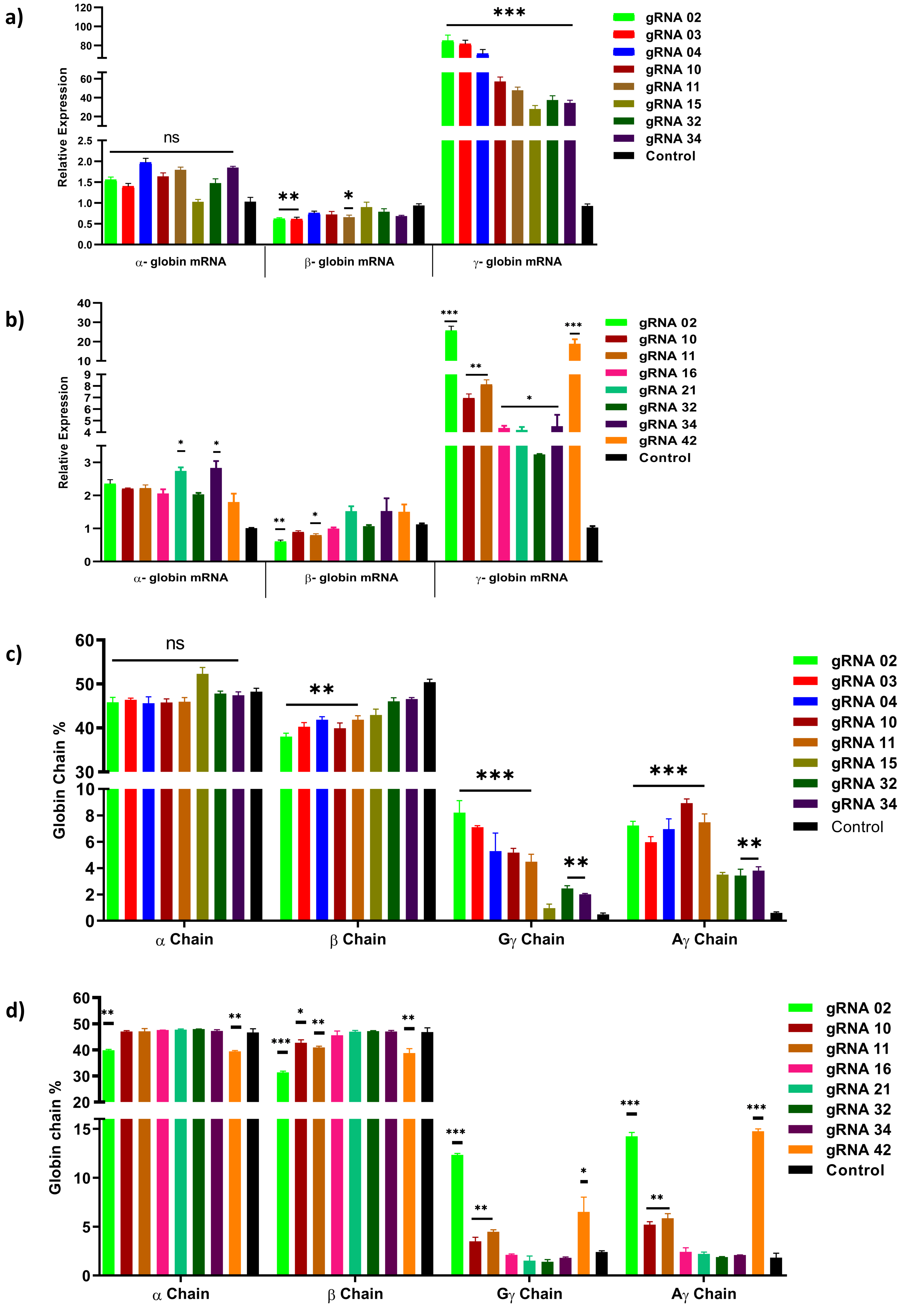


#### Supplementary Figure 7

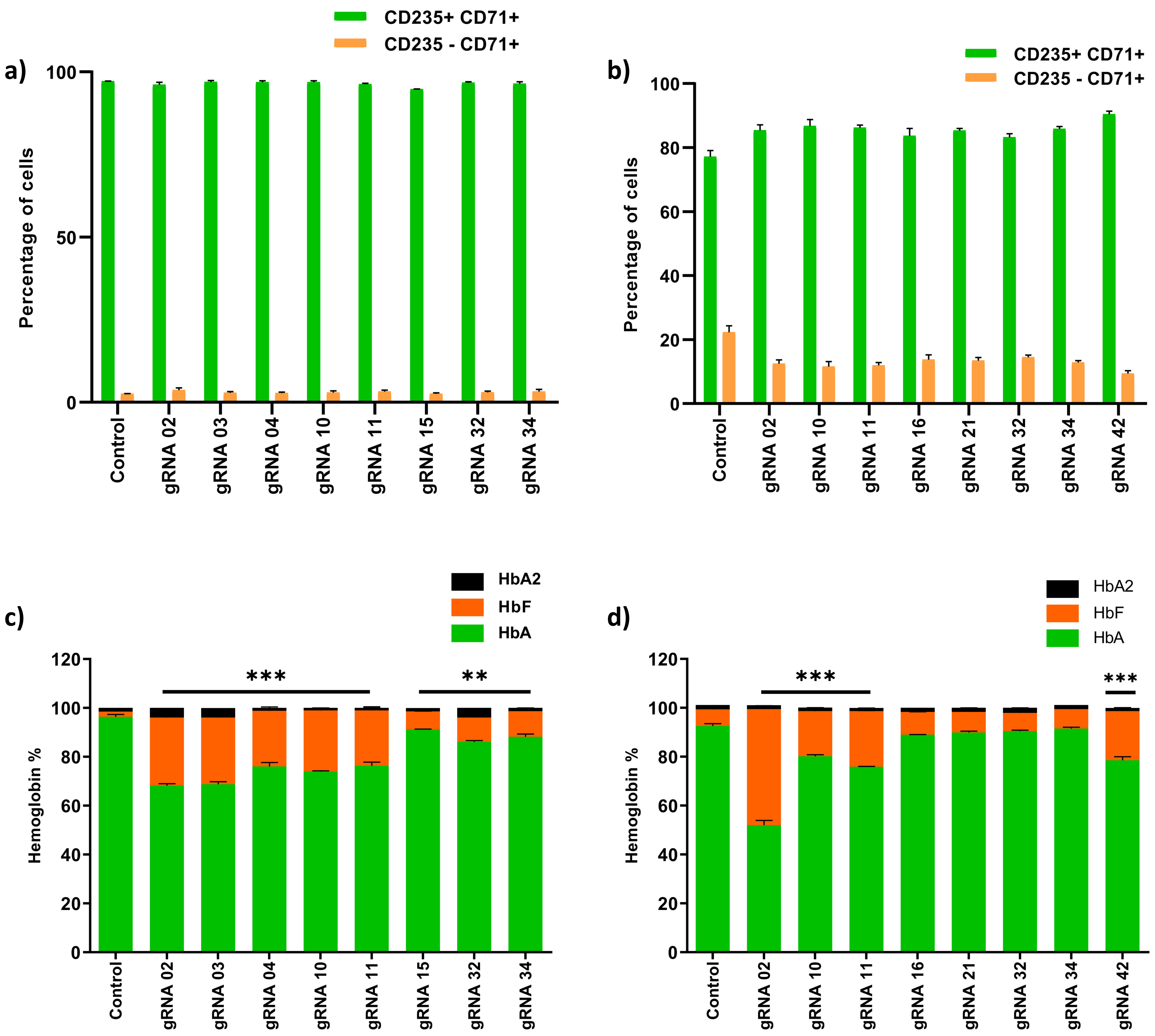


#### Supplementary Figure 8

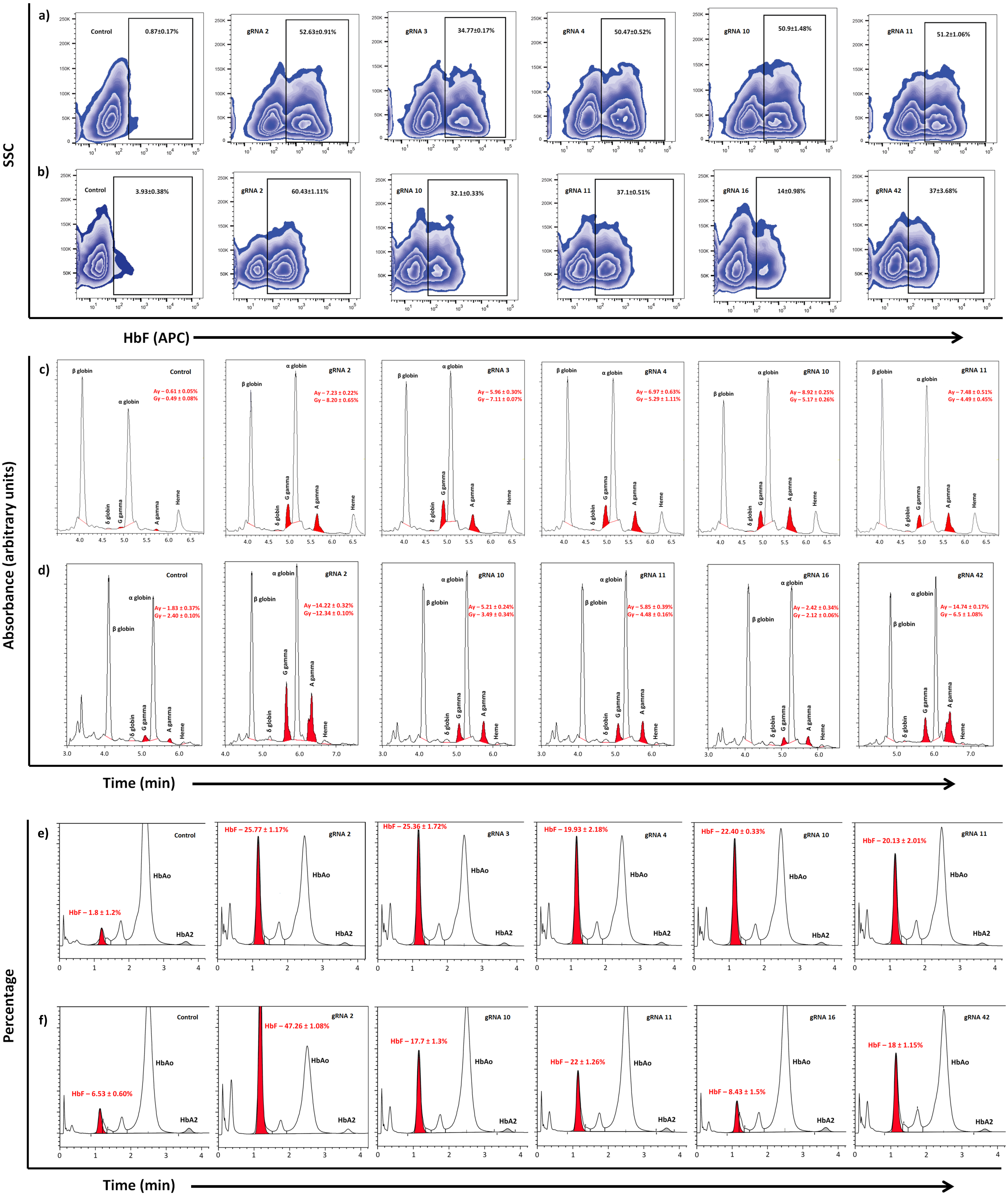


### Supplementary Figure 9

#### SUPPLEMENTAL FIGURE LEGENDS

**S1: Cumulative editing efficiency of individual gRNA at different target sites of *HBG* promoter determined using NGS.** Comparison of total editing efficiency mediated by ABE (A) and CBE (B) for different gRNAs in the target site, the data is represented as a percentage of total modified reads and total unmodified reads.

**S2: Effect of target base position on base substitution efficiency:** Role of the positional effect of a target base in deciding the editing scope of ABE (A) and CBE (B) for different gRNAs. The x-axis represents the base position of individual gRNAs, and the y-axis represents the base substitution percentage. In the case of ABE, an active editing window was observed between 3-9th bp while in CBE, the editing window was from 1-9th bp.

**S3: Effects of adenine and cytosine base editors mediated editing on product purity.** Specific and non-specific editing of ABE (A) and CBE (B) at the target sites for different gRNAs analyzed by deep sequencing (NGS); individual base position in the *HBG* promoter region is mentioned in the x-axis, and the percentage of specific and non-specific conversion is represented in the y-axis as the number of reads. There were more non-specific base conversions in CBE compared to ABE.

**S4: Indel profile of individual gRNAs:** Percentage of indels at the indicated target sites in the cells treated with ABE (A) and CBE (B) analyzed by NGS. The frequency of indels, as evident from NGS data, is very less in ABE when compared to CBE.

**S5: Analysis of base substitution by ABE and CBE through Sanger sequencing:** HUDEP2 cells expressing ABE (A) or CBE (B) transduced with different gRNAs were sequenced by Sanger sequencing and analyzed using EditR. The individual base position in the *HBG* promoter and the percentage conversion of the respective bases is represented in the x-axis and y-axis (A to G or C to T), respectively.

**S6: Total editing efficiency of individual gRNAs at different target sites of *HBG* promoter determined by deep sequencing and their corresponding indels:** Validation of cumulative editing efficiency mediated by ABE (A) and CBE (C) for the top 8 targets, the data represented as total modified reads to total unmodified reads. Analysis of indel frequency at the indicated sites in the cells treated with ABE (B) CBE (D). Data are expressed as means ± SEM from three independent biological replicates.

**S7: Robust induction of gamma-globin upon base editing the potential targets in *HBG* promoter.** Relative expression of globin genes in adenine base edited samples (A), and cytosine base edited samples (B) before differentiation by using qRT-PCR; RP-HPLC analysis of globin chains in ABE (C) and CBE (D) expressing HUDEP2 cells transduced with the individual gRNAs after differentiation. All data expressed as means ± SEM from three independent biological replicates. Asterisks indicate levels of statistical significance ***$p<0.001$; **$p<0.01$; *$p<0.05$.

**S8: Base editing of the target region in *HBG* promoter does not affect erythroid differentiation and up-regulates HbF level:** FACS analysis of CD235a and CD71 expression in erythroblast derived from HUDEP2 cells expressing ABE (A) or CBE (B) transduced with indicated gRNAs. HPLC analysis of total hemoglobin in ABE (C) and CBE (D) expressing in HUDEP2 cells transduced with the respective gRNA after erythroid differentiation. All data expressed as means ± SEM from three independent biological replicates. Asterisks indicate levels of statistical significance ***$p<0.001$; **$p<0.01$; *$p<0.05$.

**S9. Adenine base editing or cytosine base editing in the target sites at *HBG* promoter reactivates fetal globin expression in human erythroid progenitors.** HbF positive cells in HUDEP2 cells expressing ABE (A) or CBE (B) transduced with the indicated gRNAs before erythroid differentiation by flow cytometry analysis are represented as zebra plots. HPLC chromatogram profile for globin chains (C and D) and hemoglobin variants (E and F) showing the distribution for the top 5 selected candidates of ABE (Control, gRNA 2, 3, 4, 10 and 11) and CBE (Control, gRNA 2, 10, 11, 16 and 42) in sequential order. All the representative plots are shown with the mean value ± SE for the three independent biological replicates.

#### SUPPLEMENTAL TABLES

**Table 1:**

| Sl.no | Names | gRNA Info 5'-3' | Editing window for CBE | Editing window for ABE |
|---|---|---|---|---|
| **1** | gRNA 1 | ggctagggatgaagaataaa | yes | yes |
| **2** | gRNA 2 | cttgaccaatagccttgaca | yes | yes |
| **3** | gRNA 3 | atAtttgcattgagatagtg | yes | yes |
| **4** | gRNA 4 | gtggggaaggggcccccaag | nil | yes |
| **5** | gRNA 5 | tggtcaagtttgccttgtca | yes | yes |
| **6** | gRNA 6 | gtttgccttgtcaaggctat | yes | nil |
| **7** | gRNA 7 | cttgtcaaggctattggtca | yes | yes |
| **8** | gRNA 8 | caaggctattggtcaaggca | yes | yes |
| **9** | gRNA 9 | gctattggtcaaggcaaggc | nil | yes |
| **10** | gRNA 10 | aggcaaggctggccaaccca | yes | yes |
| **11** | gRNA 11 | ggcaaggctggccaacccat | yes | yes |
| **12** | gRNA 12 | aaggctggccaacccatggg | yes | nil |
| **13** | gRNA 13 | cccatgggtggagtttagcc | yes | yes |
| **14** | gRNA 14 | ccatgggtggagtttagcca | nil | yes |
| **15** | gRNA 15 | gctaaactccacccatgggt | yes | yes |
| **16** | gRNA 16 | ccctggctaaactccaccca | yes | yes |
| **17** | gRNA 17 | tatctgtctgaaacggtccc | yes | nil |
| **18** | gRNA 18 | tatttgcattgagatagtgt | yes | yes |
| **19** | gRNA 19 | atgcaaatatctgtctgaaa | yes | Yes |

| | | | | |
|---|---|---|---|---|
| 20 | gRNA 20 | ggaatgactgaatcggaaca | yes | yes |
| **21** | gRNA 21 | actgaatcggaacaaggcaa | yes | yes |
| **22** | gRNA 22 | aaaaactggaatgactgaat | yes | yes |
| **23** | gRNA 24 | gcattgagatagtgtgggga | nil | yes |
| **24** | gRNA 25 | attgagatagtgtggggaag | nil | yes |
| **25** | gRNA 29 | agaataaattagagaaaaac | nil | yes |
| **26** | gRNA 30 | ggagaaggaaactagctaaa | nil | yes |
| **27** | gRNA 32 | cagttccacacactcgcttc | yes | yes |
| **28** | gRNA 33 | cttcatccctagccagccgc | yes | yes |
| **29** | gRNA 34 | cctagccagccgccggcccc | yes | yes |
| **30** | gRNA 35 | ccgccggcccctggcctcac | yes | nil |
| **31** | gRNA 36 | actggatactctaagactat | yes | yes |
| **32** | gRNA 37 | ccaggggccggcggctggct | yes | yes |
| **33** | gRNA 38 | tgaggccaggggccggcggc | yes | yes |
| **34** | gRNA 39 | ttagagtatccagtgaggcc | nil | yes |
| **35** | gRNA 40 | tagtcttagagtatccagtg | yes | yes |
| **36** | gRNA 41 | tagagtatccagtgaggcca | yes | yes |
| **37** | gRNA 42 | agagtatccagtgaggccag | yes | yes |
| **38** | gRNA 43 | ccagtgaggccaggggccgg | nil | yes |
| **39** | gRNA 44 | caggggccggcggctggcta | yes | nil |
| **40** | gRNA 45 | aagcagcagtatcctcttgg | yes | yes |
| **41** | gRNA 46 | attaagcagcagtatcctct | yes | yes |
| **42** | control | Plasmid without gRNA | - | - |

**Table 2:**

| Sl. No | Primer name | Primer sequence 5'- 3' | Band size | Source |
|---|---|---|---|---|
| **1** | Sequencing F | gagggcctatttcccatgat | 539bp | This study |
| **2** | Sequencing R | tggatctctgctgtccctgt | | This study |
| **3** | HBF 1 F | acaaaagaagtcctggtatc | 490bp | This study |
| **4** | HBF 2 F | ttactgcgctgaaactgtgg | 772bp | This study |
| **5** | HBF 1 R | cttcccagggtttctcctcc | | This study |
| **6** | HBB E2E3 RT F | acctttgccacactgagtgag | 110bp | This study |
| **7** | HBB E2E3 RT R | tttgccaaagtgatgggcca | | This study |
| **8** | HBA RT F | cgacaagaccaacgtcaagg | 99bp | This study |
| **9** | HBA RT R | gtggggaaggacaggaacat | | This study |
| **10** | HBF E2E3 RT F | cttccttgggagatgccata | 136bp | This study |
| **11** | HBF E2E3 RT R | aaaacggtcaccagcacatt | | This study |
| **12** | GAPDH E7E8 RT F | ctgcaccaccaactgcttag | 110bp | This study |
| **13** | GAPDH E7E8 RT R | gtcttctgggtggcagtgat | | This study |
| **14** | NGS 2 F | gctcttccgatct tgaatcggaacaaggcaaagg | 325bp | This study |
| **15** | NGS 2 R | gctcttccgatct gtgaaatgacccatggcgtc | | This study |
| **16** | NGS 3 F | gctcttccgatct cctggacctatgcctaaaaca | 318bp | This study |
| **17** | NGS 3 R | gctcttccgatct agtttagccagggaccgttt | | This study |
| **18** | NGS 4 F | gctcttccgatct cggctgacaaaagaagtcct | | This study |
| **19** | HBG1 F | ccacagtacctgccaaagaa | 940bp | This study |
| **20** | HBG2 F | ccatagtatctggtaaagagca | 940bp | This study |
| **21** | HBG1/2 R | ggcgtctggactaggag | | This study |

**Table 3:**

| **Sl.no** | **Names** | **Primers used for Sanger sequencing** | **Primers used for NGS** | **Primers used for NGS HBG1/HBG2** |
|---|---|---|---|---|
| **1** | gRNA 1 | HBF 1 F/HBF1R | NGS 2 F/R | Not applicable |
| **2** | gRNA 2 | HBF 1 F/HBF1R | NGS 2 F/R | NGS4 F/NGS 2 R |
| **3** | gRNA 3 | HBF 1 F/HBF1R | NGS 2 F/R | NGS4 F/NGS 2 R |
| **4** | gRNA 4 | HBF 1 F/HBF1R | NGS 2 F/R | NGS4 F/NGS 2 R |
| **5** | gRNA 5 | HBF 1 F/HBF1R | NGS 2 F/R | Not applicable |
| **6** | gRNA 6 | HBF 1 F/HBF1R | NGS 2 F/R | Not applicable |
| **7** | gRNA 7 | HBF 1 F/HBF1R | NGS 2 F/R | Not applicable |
| **8** | gRNA 8 | HBF 1 F/HBF1R | NGS 2 F/R | Not applicable |
| **9** | gRNA 9 | HBF 1 F/HBF1R | NGS 2 F/R | Not applicable |
| **10** | gRNA 10 | HBF 1 F/HBF1R | NGS 2 F/R | NGS4 F/NGS 2 R |
| **11** | gRNA 11 | HBF 1 F/HBF1R | NGS 2 F/R | NGS4 F/NGS 2 R |
| **12** | gRNA 12 | HBF 1 F/HBF1R | NGS 2 F/R | Not applicable |
| **13** | gRNA 13 | HBF 1 F/HBF1R | NGS 2 F/R | NGS4 F/NGS 2 R |
| **14** | gRNA 14 | HBF 1 F/HBF1R | NGS 2 F/R | NGS4 F/NGS 2 R |
| **15** | gRNA 15 | HBF 1 F/HBF1R | NGS 2 F/R | NGS4 F/NGS 2 R |
| **16** | gRNA 16 | HBF 1 F/HBF1R | NGS 2 F/R | NGS4 F/NGS 2 R |
| **17** | gRNA 17 | HBF 1 F/HBF1R | NGS 2 F/R | Not applicable |
| **18** | gRNA 18 | HBF 1 F/HBF1R | NGS 2 F/R | Not applicable |
| **19** | gRNA 19 | HBF 1 F/HBF1R | NGS 2 F/R | NGS4 F/NGS 2 R |
| **20** | gRNA 20 | HBF 2 F/HBF1R | NGS 3 F/R | NGS4 F/NGS 2 R |

| | | | | |
|---|---|---|---|---|
| **21** | **gRNA 21** | **HBF 2 F/HBF1R** | **NGS 3 F/R** | **NGS4 F/NGS 2 R** |
| **22** | gRNA 22 | HBF 2 F/HBF1R | NGS 3 F/R | Not applicable |
| **23** | gRNA 24 | HBF 2 F/HBF1R | NGS 2 F/R | Not applicable |
| **24** | gRNA 25 | HBF 2 F/HBF1R | NGS 2 F/R | Not applicable |
| **25** | gRNA 29 | HBF 2 F/HBF1R | NGS 3 F/R | Not applicable |
| **26** | gRNA 30 | HBF 2 F/HBF1R | NGS 3 F/R | NGS4 F/NGS 2 R |
| **27** | gRNA 32 | HBF 1 F/HBF1R | NGS 2 F/R | NGS4 F/NGS 2 R |
| **28** | gRNA 33 | HBF 1 F/HBF1R | NGS 2 F/R | Not applicable |
| **29** | gRNA 34 | HBF 1 F/HBF1R | NGS 2 F/R | NGS4 F/NGS 2 R |
| **30** | gRNA 35 | HBF 1 F/HBF1R | NGS 2 F/R | Not applicable |
| **31** | gRNA 36 | HBF 1 F/HBF1R | NGS 2 F/R | NGS4 F/NGS 2 R |
| **32** | gRNA 37 | HBF 1 F/HBF1R | NGS 2 F/R | Not applicable |
| **33** | gRNA 38 | HBF 1 F/HBF1R | NGS 2 F/R | Not applicable |
| **34** | gRNA 39 | HBF 1 F/HBF1R | NGS 2 F/R | Not applicable |
| **35** | gRNA 40 | HBF 1 F/HBF1R | NGS 2 F/R | NGS4 F/NGS 2 R |
| **36** | gRNA 41 | HBF 1 F/HBF1R | NGS 2 F/R | Not applicable |
| **37** | gRNA 42 | HBF 1 F/HBF1R | NGS 2 F/R | NGS4 F/NGS 2 R |
| **38** | gRNA 43 | HBF 1 F/HBF1R | NGS 2 F/R | Not applicable |
| **39** | gRNA 44 | HBF 1 F/HBF1R | NGS 2 F/R | NGS4 F/NGS 2 R |
| **40** | gRNA 45 | HBF 2 F/HBF1R | NGS 3 F/R | Not applicable |
| **41** | gRNA 46 | HBF 2 F/HBF1R | NGS 3 F/R | Not applicable |
| **42** | control | HBF 1 F, 2 F/HBF1R | NGS 2 and NGS 3 | NGS4 F/NGS 2 R |
